# Deciphering the role of autophagy under Cd toxicity in *Arabidopsis thaliana*

**DOI:** 10.1101/2025.08.27.672299

**Authors:** Aurelio M. Collado-Arenal, Felipe L. Pérez-Gordillo, Jesús Espinosa, Rubén Moreno-Díaz, Sergey Shabala, María C. Romero-Puertas, Luisa M. Sandalio

**Affiliations:** Estación Experimental del Zaidín, Department of Stress, Development and Signalling in Plants. Consejo Superior de Investigaciones Científicas (CSIC); School of Biological Science and ARC Training Centre for Smart & Sustainable Horticulture, University of Western Australia, Perth WA6009 Australia; Internatiolal Research Centre for Environmental Membrane Biology, Foshan University, Foshan, China

**Keywords:** Autophagy, cadmium, ionomic, oxidative stress, transporters

## Abstract

Cadmium (Cd) is a toxic pollutant in soil and water affecting plants, animals, and humans. Autophagy, a cellular recycling process, is crucial for different biotic and abiotic plant stress responses. This study explores the autophagy role in *Arabidopsis* under Cd stress, using wild-type, autophagy-deficient mutants (*atg5*, and *atg7*) and overexpressing lines (*35S:ATG5*, *35S:ATG7)*. Cd exposure induced autophagy, as evidenced by ATG8a and ATG8a-PE accumulation, GFP-ATG8a fluorescence, and upregulation of ATG genes and proteins. Responses differed between Col-0 and Ws backgrounds, with Ws showing higher Cd tolerance. *atg5* mutants were more sensitive to Cd, indicating the autophagy protective role, whereas *ATG5/ATG7* overexpression did not significantly enhance Cd tolerance. Although oxidative stress may activate autophagy, *ATG5/ATG7* overexpression did not significantly change the oxidative stress response. Notably, *atg5* mutants displayed marked disruptions in metal/ion homeostasis under control and Cd conditions, reinforcing autophagy’s role inion homeostasis. In contrast, *atg7* showed no significant differences from its WT (Ws), suggesting genotype-specific effects. Transcriptional analysis of metal transporters and ion flux analyses indicates that autophagy can regulate metal/ion accumulation through transcriptional control and post-translational modifications (e.g., ROS) with a differential response being observed between Ws and Col-0 plants.

## Introduction

Cadmium is a toxic heavy metal for plant, animals and humans which is accumulated in the soil as a consequence of industrial contamination, metal smelters, batteries, pigment manufacturing, plastic production, burning fossil fuels and the incineration of municipal waste, as well as mines and phosphate fertilizers (Peco et al., 2020; Sidhu and Bali, 2022). Currently, between 14% and 17% of surface soils exceed agricultural thresholds for heavy metal contamination, corresponding to 242 million hectares, or 16% of the world’s cropland. In particular, the global excess rate of Cd, which is a non-essential metal, is the highest, reaching 9% (Hou et al., 2025). Unfortunately, the issue of soil contamination is expected to continue to increase due to the growth in demand for critical metals needed for the “green transition” of zero net emissions and the development of wind turbines, electric vehicle batteries, and photovoltaic devices (Hou et al., 2025). Being accumulated in plants, Cd enters the food chain, leading to significant risks for human health and threatening water quality and food security. High Cd concentration also restricts plant growth and yield by affecting a broad range of physiological and metabolic processes (Sandalio et al., 2001; Noguerol et al., 2016; Carvalho et al., 2020; Peco et al., 2020).

One of the primary consequences of Cd stress in plants is the over production of reactive oxygen species (ROS) such as H_2_O_2_,**·**OH, O_2_^.-^, and ^1^O_2_ (Collado-Arenal et al., 2024; Cuypers et al., 2023; Rodríguez-Serrano et al., 2009). While the effect is indirect (Cuypers et al., 2023; Sandalio et al., 2012), Cd accumulation triggers oxidative damages to most macromolecules, lipids, proteins and DNA (Sandalio et al., 2012). Under oxidative conditions oxidized proteins, obsolete proteins or damaged/obsolete organelles must be removed to avoid further and worse consequences. This process can be regulated by autophagy which is a mechanism of degradation and recycling of unnecessary or dysfunctional cellular components (including organelles), which takes place in all eukaryotes, and it is of extreme importance for cell survival (Avin-Wittenberg et al., 2018; Petersen et al., 2024a). This process requires the coordinated action of approximately 40 genes which are evolutionary highly conserved and grouped under the term autophagy related genes (ATGs; (Avin-Wittenberg et al., 2018; Petersen et al., 2024b). The different ATGs integrate different signals resulting in the formation of autophagosomes which are further degraded in the vacuole (Avin-Wittenberg et al., 2018; Petersen et al., 2024a). ATGs are grouped into different complexes. This includes: (1) the ATG1 protein kinase complex (containing ATG13, ATG101; ATG11 and ATG1), involved in controlling the initiation of autophagy; (2) the ATG9 complex, linked to vesicles, which is recruited to the pre-autophagosomal structure (PAS) promoting the initial expansion of the isolation membrane; (3) the VPS34/PI3K complex (including ATG6, ATG14, VPS34 and VPS1), which produces phosphatidylinositol 3-phosphate (PI3P) on the isolation membrane; (4) the ATG2–ATG18 complex, which regulates the expansion of the isolation membrane and two ubiquitination-like conjugation systems; (5) the ATG12–ATG5 conjugation system that includes ATG16, ATG5, ATG12, ATG10 and ATG7; and (6) the ATG8 lipidation system, which insert ATG8 phosphatidylethanolamine (PE) into the expanding isolation membrane thus promoting the elongation of the isolation membrane (Avin-Wittenberg et al., 2018; Stephani and Dagdas, 2020; Yagyu and Yoshimoto, 2024). These functional complexes work together to orchestrate the formation of autophagosomes.

The ATG8 lipidation system requires a previous post-translational processing of ATG8 by cysteine protease ATG4 (Avin-Wittenberg et al., 2018; Stephani and Dagdas, 2020; Yagyu and Yoshimoto, 2024). ATG8–PE provides a docking platform for macroautophagy adaptors to facilitate autophagosome formation, or for receptor recognition to selectively recruit cargos to the isolation membrane. Several ATG8 isoforms have been identified in plants and play an important role in autophagosome formation and functioning in all the plant species studied (Rogov et al., 2023) and the different isoforms could bind distinct sets of plant proteins with varying degrees of overlap (Zess et al., 2019). Lipidated ATG8 interacts with cargo receptors to select unwanted material (proteins and organelles) which are then delivered to the vacuole for degradation (Avin-Wittenberg et al., 2018; Petersen et al., 2024a). Another component of autophagy is the target of rapamycin (TOR) which is a highly conserved protein kinase that serves as a central regulator of growth and metabolism in both plants and animals. TOR integrates signals from nutrients, energy levels, hormones, growth factors, and environmental cues to coordinate cellular responses (Dong et al., 2022; Shi et al., 2018) and TORC1 was shown to operate as a negative regulator of autophagy (Díaz-Troya et al., 2008). The TOR complex is linked to the modulation of the ATG1 complex in response to different stress conditions (Mugume et al., 2020).

Autophagy is involved in the adaptation of plants to a wide range of environmental stresses including nutrient starvation, heat stress, drought, salt, and pathogen invasion (Hasan et al., 2017; Liu et al., 2022; Thirumalaikumar et al., 2021; Zhou et al., 2015; Petersen et al., 2024a). However, the role of autophagy in response to heavy metals in plants remains elusive (Hasan et al., 2017), although upregulation of different ATGs in plants exposed to some metals have been reported (Zhou et al., 2015). It was shown also that Cd induces pexophagy, a selective degradation of peroxisomes, in response to short period of treatment as a mechanism to balance the number of peroxisomes and their quality favoring degradation of oxidative modified peroxisomes, thus regulating redox homeostasis in the cell (Calero-Muñoz et al., 2019). However, the information available on the regulation of autophagy by Cd and the role of this process in the cell response to Cd is very scarce. Several studies suggested that ROS produced under abiotic and biotic stress can activate SnRK2 which in its turn activates autophagy by inhibiting TOR kinase (Rosenberger and Chen, 2018; Signorelli et al., 2019). Additionally, redox regulation of ATG4 and ATG3 have been also reported in Chlamydomones and plants (Mallén-Ponce and Pérez-Pérez, 2023; Pérez-Martín et al., 2014). However, ATG4 oxidation can also lead to ATG4 inactivation through oligomer formation in yeasts and algae (Pérez-Pérez et al., 2021, 2016). Therefore, understanding the mechanisms involved in autophagy regulation and its role in the mechanism of the tolerance of plants to heavy metals is a very important issue for food security, phytoremediation and plant biodiversity research.

In this study, we attempted to fill in the above knowledge gaps by comparing autophagy-related processes in two *Arabidopsis thaliana* ecotypes with various Cd tolerance (Col-0 – sensitive; Wassileskija (Ws) – tolerant; (Amaral dos Reis et al., 2021) and their KO and overexpressing lines. Our results demonstrate that *atg5* mutants displayed higher oxidative stress and marked disruptions in metal/ion homeostasis under control and Cd conditions, reinforcing autophagy’s role in nutrient balance and detoxification. In contrast, *atg7* showed no significant differences from its WT (Ws), suggesting genotype-specific effects. Transcriptional analysis of metal transporters indicates that autophagy regulates metal/ion accumulation through transcriptional control and post-translational modifications (e.g., ROS, NO, phosphorylation).

## Material and Methods

*Arabidopsis thaliana* ecotype Columbia-0 (Col-0) constitutes the genetic background for all wild-type mutants plants used in this study, except for *atg7* mutants, which were obtained in a Wassilewskija (Ws) background. Wild-type plants expressing green fluorescent protein (GFP)-ATG8a and Arabidopsis T-DNA disruption mutants affecting *atg5* and *atg7* and the over-expresser *35S:ATG5 and 35S:ATG7* were kindly supplied by Dr. Vierstra (Washington University, St Louis, USA). Seeds were sown in cut Eppendorf tubes, containing semisolid ½ strength Hoagland nutrient solution (Hoagland and Arnon, 1950) in hydroponic cultures and the plantlets were grown in a growth chamber at 22 ± 1 °C; day/night 16/8 h; irradiance 120–150 μM m^-2^ s^-1^. Afterwards, plants were exposed to 50 μM CdCl_2_ treatment and analyzed at various time points, from 1 to 10 d after Cd treatment. Alternatively, Arabidopsis plants were grown in vertical plates containing Murashige & Skoog (MS) 0.5X solid medium (Murashige and Skoog, 1962) containing 3% sucrose (w/v) and 0.8%, phytoagar (w/v) ± 25 μM CdCl_2_. Seeds were surface sterilized and stratified for 48 h at 4 °C, sown on MS plates, and grown as described above.

### Growth parameters data and photosynthetic pigments content

In plants grown in vertical plates, pictures were taken at 12 days and root length analyzed by ImageJ software. In adult plants grown under hydroponic conditions basic agronomical characteristics such as leaf and root fresh weight, number and area of leaves, root length and rosette area, were analyzed. Pigment composition (chlorophyll and carotenoid contents) were analyzed in fresh leaves after extraction in 80% acetone and quantified spectrophotometrically following the method of Lichtenthaler and Wellburn, (1983) (Lichtenthaler and Wellburn, 1983).

### Plant extracts, electrophoresis and Western blot analysis

Whole ATG8a-GFP plants (100–200 mg) were homogenized in a 100 mM Tris-HCl, pH 7.5, 400 mM sucrose, 1 mM ethylenediaminetetra-acetic acid (EDTA), 10 mg ml^−1^ of sodium deoxycholate, 0.1 mM phenylmethylsulfonyl fluoride, 10 mg ml^−1^ of pepstatin A, and 4 % (v/v) protease inhibitor cocktail (Roche) centrifuged at 14.000 rpm for 30 min to obtain the supernatant fraction as described previously (Calero-Muñoz et al., 2019). For immunoblot analysis, 40 μg of leaf protein extracts was separated on 12-15 % acrylamide gels. The proteins were then transferred to a polyvinylidene fluoride membrane (Millipore Co., Bedford, MA, USA) in a Bio-Rad Semi-Dry Transfer Cell (Bio-Rad, Hercules, CA, USA). The membranes were incubated for 1 h in blocking buffer containing 5 % milk powder (w/v) prepared in Tris Buffer Saline (TBS) containing 0.1 % Tween 20 (v/v). Antibodies were diluted in blocking buffer: anti-ATG8 (AS14 2769, Agrisera, Sweden; 1:2.000) and anti-GFP (AS20 4443, Agrisera, Sweden; 1:1.000). Membranes were washed with TBS and incubated with goat anti-rabbit IgG conjugated with horseradish peroxidase (HRP) (AS09 602, Agrisera, Sweden; 1:10.000 in blocking buffer). After washing with TBS, bands were visualized by the ECL Western blot detection system Amersham ECL Western Blotting Detection Kit. The immunodetected protein bands were quantified relative to total proteins by imaging with a Stain Free method using 2,2,2-tricloroethanol (TCE) and activated in gel by UV illumination for 1 min. Protein bands were quantified using ImageJ and ImageLab software.

Plant material was homogenized as described above and Western blot analysis were carried out using different antibodies: anti-ATG5 (AS15 3060, Agrisera, Sweden; 1:2.000), anti-ATG8 (AS14 2769, Agrisera, Sweden; 1:2.000), anti-NBR1 (AS14 2805, Agrisera, Sweden; 1:1.000) and anti-RPT5A (BML-PW8770, Enzo, EEUU; 1:2.000).

### Autophagy flux analysis

Autophagy flux analysis was carried out by analysing ATG8 and ATG8-lipidated by SDS-PAGE and further Western blot analysis using antibodies against ATG8a (Calero-Muñoz et al., 2019). Alternatively, the analysis of free and attached GFP to ATG8a was analyzed by using Arabidopsis lines expressing the protein fusion ATG8a-GFP (supplied by Dr. Vierstra). After the treatment, plant extracts were obtained and proteins separated by SDS-PAGE and further Western blot using GFP antibodies as mentioned previously. Additionally, ATG8a-GFP plants were used to analyse GFP fluorescence in leaf discs in a multiplate using a fluorimeter ClarioStar (BMG biotech, Cary, NC) measuring at 488 ± 8 nm of excitation and 530 ± 8 nm of emission.

### Enzyme extractions and assays

All operations were carried out at 4 °C. Frozen tissues (500 mg fresh weight) were ground in a mortar with liquid nitrogen and 1 ml of buffer (50 mM Tris-HCl pH 7.5), containing 0.1 mM EDTA, 0.2% (v/v) Triton X-100, 2 mM DTT and protease inhibitor cocktail (Sigma). The homogenate was centrifuged at 14,000 rpm for 30 min and the supernatants were collected and used for different enzyme assays. Catalase (CAT, EC 1.11.1.6) and guaiacol peroxidase (GPX, EC 1.11.1.7) activities were analysed according to Peco et al., (2020) (Peco et al., 2020) and glycolate oxidase (GOX, EC 1.1.3.15) as Kerr and Groves (1975) (Kerr and Groves, 1975).

Protein content was determined by the Bradford method (1976) (Bradford, 1976) using bovine serum albumin (BSA) as standard.

### Mineral analysis

Plant samples were oven-dried at 60 °C for 3 d. Roots and leaves were weighed to determine the dry mass and were digested with an HNO_3_/H_2_O_2_ mixture using a microwave digestion system (ETHOS 1, Milestone). Mineral composition was measured by inductively coupled plasma optical emission spectrometry (ICP-OES, Varian 720-ES, CEBAS-CSIC, Murcia).

### Oxidative stress markers

Lipid peroxidation was analyzed on the basis of malondialdehyde (MDA) concentration as described by Buege and Aust (1978) (Buege and Aust, 1978). H_2_O_2_ concentration was analyzed by a spectrofluorometric method using homovanilic acid as described by Romero-Puertas et al. (2004) (Romero-Puertas et al., 2004).

### Histochemical localization of H_2_O_2_ and O ^.-^

H_2_O_2_ and O_2_^.-^ accumulation were imaged in leaves and roots by histochemical staining according to Romero-Puertas et al., (2004) (Romero-Puertas et al., 2004). For H_2_O_2_ imaging the tissues were immersed in a 0.1 % (w/v) solution of DAB (3,3’-diaminobenzidine), vacuum-infiltrated for 5 min and finally incubated in the dark at room temperature overnight. For O ^.-^ staining, leaves and roots were immersed in a 0.1 % (w/v) solution of Nitro Blue Tetrazolium (NBT) containing 10 mM Na-azide and were vacuum infiltrated for 10 min and illuminated until dark blue spots appeared. To improve imaging spots, samples were incubated in ethanol 96 % overnight and further boiling for 5 min.

### Confocal microscopic analyses

To image autophagosomes, Arabidopsis GFP-ATG8a lines were used and GFP was visualized after 7 d days of Cd treatment using a confocal Leyca microscope (Leica TCS SL; Leica Microsystems). To avoid the loss of GFP fluorescence during confocal microscopy analysis, the plants were incubated in liquid MS medium (with 3% sucrose) containing 0.5 μM concanamycin A (Santa Cruz Biotechnology) for 10 h at room temperature in the dark before confocal analyses. Arabidopsis plants expressing GFP-ATG8a (Exc/Em: 488/508 nm to GFP and Exc/Em 633/680 nm to chlorophylls) images were obtained as 14 z steps at 1 μm intervals by confocal microscopy, using maximum intensity projection with an optimal pinhole of 1. Analysis was carried out in leaves and roots using ImageJ software.

### qRT-PCR analysis of gene expression

Total RNA was isolated from leaves and roots by Trizol TM Reagent protocol (Thermo Fisher Scientific). RNA was treated with DNAase Kit (Invitrogen). To assess the quantity and quality of RNA a Nanodrop mySpec VWR^®^ was used and only the samples with A260/A280 ratio of 1.8-2 were used. To check the integrity of RNA, 1% agarose electrophoresis gel was carried out. For cDNA synthesis, 1 µg of total RNA and the PrimeScript RT reagent Kit (TaKaRa) were used according to the manufacturer’s protocol. qPCR was performed in a MicroAmp^®^ Fast 96-Well Reaction Plate 0.1 mL (life technologiesTM) with SYBR Green I Mix (TaKaRa) in a final volume of 10 µL containing 5 µL of SYBR Grenn I Mix, 5 µL of diluted cDNA with dH_2_O and primers 0.4 µM in a QuantStudio 3 (Applied BiosystemsTM). Each value represents three biological replicates. To analyze the amplification efficiency a standard curve of four serial dilutions points was used and the efficiency was calculated by the equation E = [10 (1/slope)-1]x100. *ATG4a*, *ATG4b*, *ATG5*, *ATG7*, *ATG8a*, *ATG8h*, *TOR*, *HMA2*, *HMA4*, *IRT1*, *IRT3*, *YSL3*, *Nramp3*, *Nramp6*, *HAK5* and *GORK* gene expression were compared with *TUB4* gene. The *TUB4* gene was selected from normalization by Gray Norm algorithm from five candidate reference genes (Terrón-Camero et al., 2020). The primers used in this work are described in Suppl. Table S1.

### Ion flux analyses

Net fluxes of Cd^2+^ were measured from the elongation zone of 10 day-old root seedlings under control conditions and in response to Cd using non-invasive microelectrode MIFE system (Shabala and Newman, 1999). Plants were immobilized in the measuring chamber 1 h before starting the experiments. Net ions fluxes were measured for 4–5 min to ensure the stability of initial ion fluxes. Cadmium chloride (CdCl_2_) at a final concentration of 50 μM was added to the bath of BSM solution (containing 500 µM KCl and 100 µM CaCl_2_) and ions fluxes were recorded for another 15 min (Hafsi et al., 2022).

### Statistical analyses

The experiments were repeated at least twice using at least three replicates. Statistical analyses were carried out by a Student’s t-test in IBM SPSS Statistics 24. Histogram figures represent media ± SEM and values followed by different letters are statistically significant at p< 0.05.

A heat map was generated using a correlation-based distance and Ward’s method for hierarchical clustering.

## Results

### Cadmium effect on autophagy process

Several approaches were used to analyzed the impact of Cd on autophagy. First, we analyzed GFP fluorescence in leaves Arabidopsis lines expressing the ATG8a-GFP at different times of treatment (0-10 d) and different CdCl_2_ concentrations (0-50 µM). The increase of GFP-dependent fluorescence was time-and concentration-dependent, with significant differences observed after 3 d of treatment (Suppl. Fig. S1A). The increase of ATG8 and lipidated ATG8a (ATG8-PE) band, as a marker of autophagy flux, was also obvious after 3 days of Cd treatment (50 µM), mainly in roots (Suppl. Fig. S1B). Additionally, we analysed autophagy by imaging GFP fluorescence in plants expressing the ATG8a-GFP by confocal microscopy using concanamycine to avoid the transport of phagosomes to the vacuole and further degradation. Under control conditions, scarce green punctates were observed (Fig. 1 A, I-III), meanwhile several autophagosomes were observed in mesophyll (Fig. 1 A, IV), epidermal (Fig. 1 A, V) and root (Fig. 1 A, VI) cells after the treatment. The Western blot analyses of leaves and roots using the antibody against the GFP demonstrated an increase of free GFP with the period of treatment (Fig. 1 B); the effect was time-dependent (Fig. 1 C). Additionally, GFP fluorescence associated to ATG8a was recorded in whole seedlings over 7 days of Cd treatment (Suppl. movie S1). The results showed an increase of fluorescence associated to ATG8a with the time of treatment in both leaves and roots.

**Figure 1.**
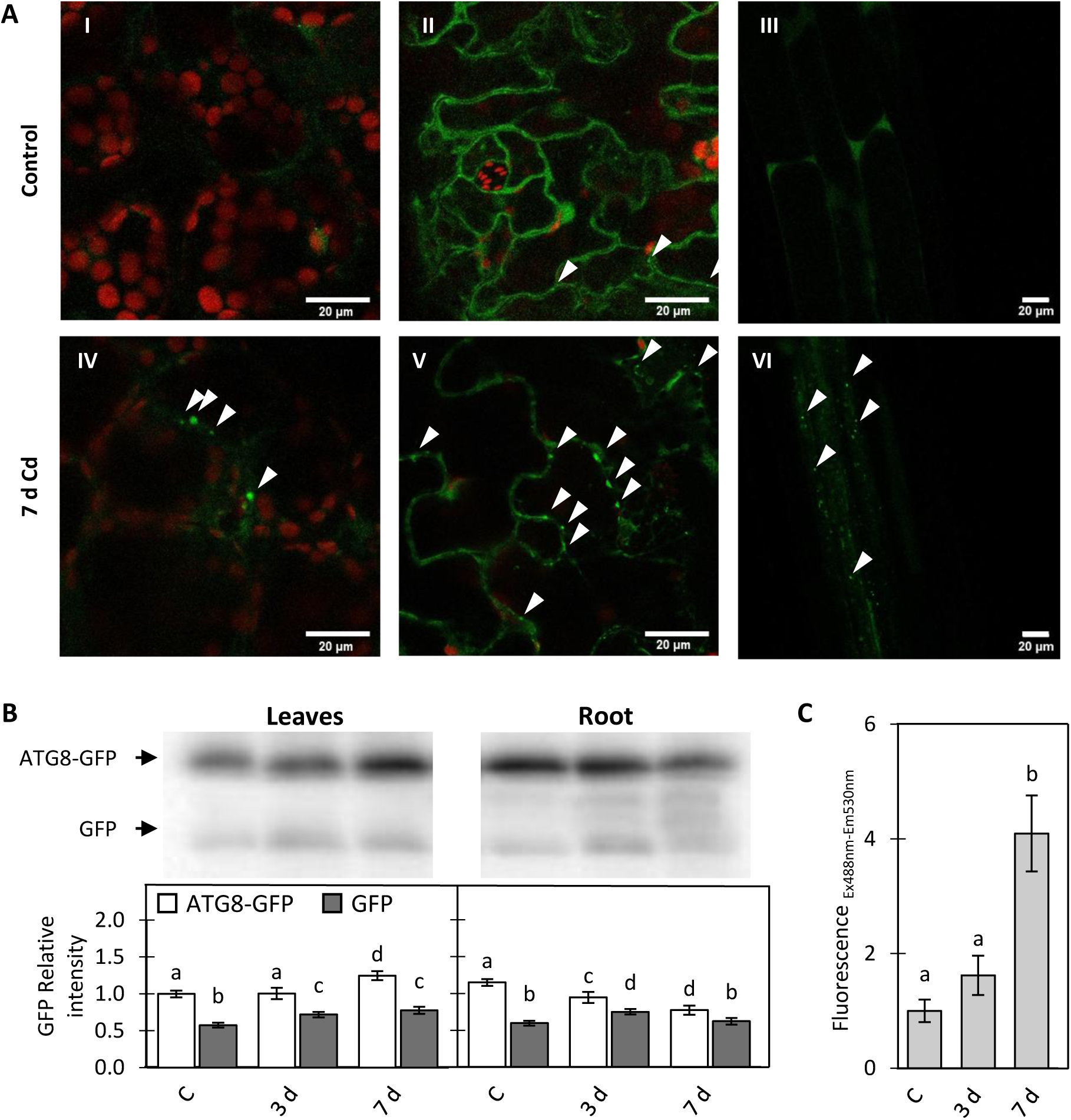
Cadmium effect on autophagy flux in Arabidopsis plants. **A**, Imaging of Cd-dependent autopghagy by confocal microscopy using Arabidopsis lines expressing the ATG8a-GFP and incubated 15 hours with concanamycine A (50 μM). Plants were grown without (control, I-III) and with 50 µM CdCl_2_ for 7 d (IV-VI). I and IV, mesophyll cells; II and V epidermal cells; III and VI root cells. Arrows indicate autophagosomes. Bars indicate 20 μm. **B**, Western Blot (40 µg of proteins/well) of leaf and root samples under control and 3 and 7 days of treatment with Cd. Membranes were incubated with α-GFP. Histogram represent the meam ± SEM of ATG8-GFP and free GFP (42 kDa and 27 kDa, respectively) in both leaves and roots at different times of Cd treatment. **C**, Fluorescence intensity (Excitation 488 nm; Emission 530 nm; mean ± SEM of 12 replicates) analysed in leaf discs from Arabidopsis lines expressing the ATG8a-GFP at different times of Cd treatment. Values followed by different letters are statistically significant at p < 0.05.

### Cadmium effect on expression of autophagy markers in wild type and autophagy disturbed Arabidopsis mutants

As autophagy markers, gene expression of *ATG4a*, *ATG4b, ATG5, ATG7, ATG8a*, *ATG8h* and *TOR* were analyzed at different times of Cd treatment in leaves and roots (Fig. 2). All lines were of Col-0 background except *atg7* (Wassilewskija (Ws) background).

**Figure 2.**
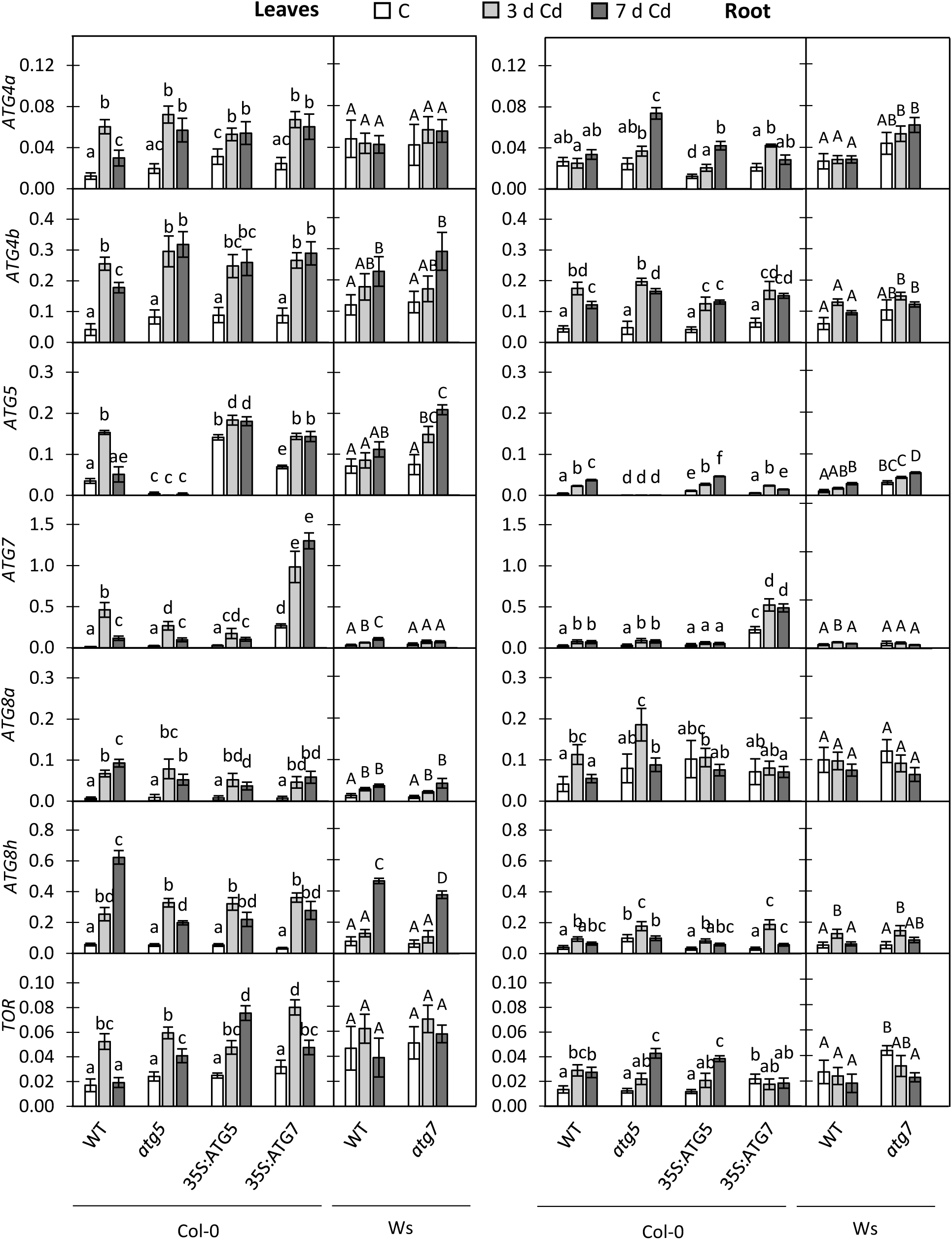
Cadmium effects on autophagy markers expression in Arabidopsis leaves and roots. Arabidopsis plants were treated with Cd (50 µM) for 3 and 7 d and the expression of autophagy markers were analysed in roots and leaves by qRT-PCR. Histograms represent mean ± SEM of 3 replicates gene expression compared with *TUB4*. Values followed by different letters are statistically significant at p < 0.05. Lowercase letters compare Col-0 backgrounds (WT, *atg5*, *35S:ATG5* and *35S:ATG7*), while capital letters compare Ws backgrounds (WT and *atg7*).

In leaves, *ATG4a* and *ATG4b* were significantly upregulated in all genotypes with Col-0 background by day 3 (Fig 2) and remaining stable after that. *ATG4a* transcript levels were much higher in leaves of lines with Ws background and not affected by Cd, while *ATG4b* genes were induced in a time-dependent manner (Fig. 2). A*TG5* was upregulated in all but *atg5* line, peaking at 3 d for WT Col-0 and at 7 d for all other lines. The highest upregulation was observed in the *35S:ATG5* over-expressor (Fig. 2). As expected, upregulation of *ATG7* was strongest in the over-expressor *35S:ATG7*; in all other lines it was much smaller and transient, peaking at 3 d. Lines with Ws background showed much smaller effects. *ATG8a* gene was upregulated in a time-dependent manner in all lines, with no effect of genetic background (e.g. Col-0 vs Ws). The same was true for *ATG8h*, although its transcript levels were 3-4 folds higher compared with *ATG8a.* Overall, the induction of ATG genes corroborate that Cd induces the autophagy processes. Additionally, we analyzed the expression of *TOR,* an autophagy regulator. A significant up-regulation of *TOR* by Cd treatment was observed in all lines with Col-0 background, peaking after 3 days (Fig 2). Plants with Ws background had constitutively higher TOR levels but unaffected by the treatment.

In roots, effects of Cd on above autophagy-related genes were generally consistent with trends reported for leaves, with a few exceptions. For example, *ATG4a* expression was not changed in WT Col-0 (Fig 2). *ATG5* levels were several-fold lower in roots compared with leaves and so were *ATG8h* and *TOR* genes while *ATG8a* transcripts were twice higher in roots.

### Accumulation of ATG8a and ATG5 in response to Cd in WT and autophagy-related mutants

Western blot analysis revealed a time-dependent increase of ATG8 and ATG8-PE in leaves of WT Col-0 plants and *atg5* mutants (Fig. 3 A). No significant changes were observed in the over-expressors *35S:ATG5* and *35S:ATG7* in both leaves and roots as expected by the induction of autophagy under control conditions (Fig. 3 A). No changes were observed in lines with Ws background, with protein levels being much lower compared to Col-0. In roots, an increase of ATG8 and ATG8-PE was observed only in WT Col-0 (Fig. 3 A). The opposite trend was observed in Ws and *atg7* plants with no significant changes being observed in leaves, while in roots a slight but significant increase of ATG8-PE was observed in WT Ws (Fig. 3 A).

**Figure 3.**
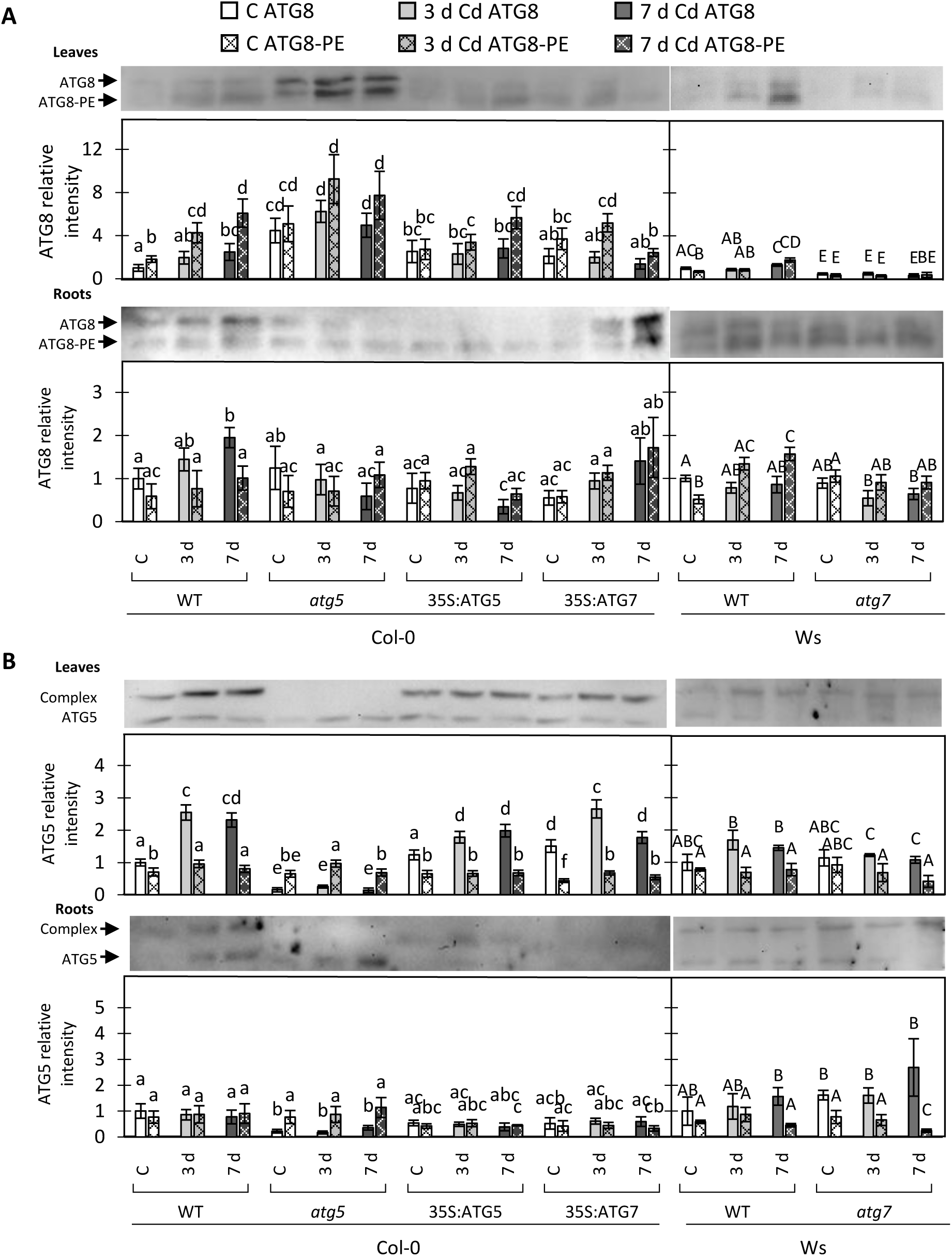
Cadmium effect on autophagy-related proteins from Arabidopsis leaves and roots after Cd treatment. Arabidopsis plants were treated with CdCl_2_ (50 µM) for 3 and 7 d and protein extract were obtained. **A**, Western blot from proteins were incubated with anti-ATG8 showing two bands: free ATG8 and ATG8 lipidated (ATG8-PE), 16 kDa and 13 kDa, respectively. **B**, Western Blot from proteins were incubated with anti-ATG5 showing two bands: ATG12 complex (ATG12-ATG16-ATG5) and free ATG5, 50 kDa and 38 kDa, respectively. Histograms represents the band intensity mean ± SEM of 3 independent replicates. Values followed by different letters (lowercase letters compare Col-0 backgrounds, while capital letters compare Ws backgrounds) are statistically significant at p < 0.05.

Free ATG5 protein slightly increased during the period of treatment in WT (Col-0) leaves, while a high significant increase was observed associated to the complex ATG12 which is an indicator of active autophagy (Fig. 3 B; (Thompson et al., 2005)). Interestingly, a thin ATG5 reactivity band was observed in *atg5*, while ATG5 associated to the complex ATG12 was absent (Fig. 3B), thus demonstrating a residual non-functional ATG5 protein. *35S:ATG5* plants did not show significant increase of free ATG5 and a slight increase in ATG5/ATG12 complex was observed. Similar results were observed in *35S:ATG7* (Fig. 3 B). Regarding Ws genotypes, free ATG5 content in leaves and roots did not change because the treatment in WT and a decrease was observed in *atg7* roots. Although not significant, a slight increase of the ATG5 associated to ATG12 complex in response to Cd was observed in both Ws lines (Fig. 3 B).

### Cadmium effect on NBR1 and proteasome markers and its regulation by autophagy

Next we analysed the accumulation of NBR1 (NEIGHBOR OF BRCA1), a selective cargo receptor in autophagy (Jung et al., 2020) in WT and ATG mutants. Cd treatment induced a time-dependent NBR1 accumulation in leaves of *atg5* genotype, consistent with the disruption of autophagy (Fig 4A). Effects were not statistically significant in other lines except Ws WT after 7 dpt. In roots, NBR1 was only accumulated at 7 dpt, mainly in WT (Col-0), followed by *atg5*, *35S:ATG5* and to a lesser extent in 35S:ATG7 (Fig. 4A). Meanwhile in Ws plants no significant changes were observed either in WT or in *atg7* (Fig. 4 A).

**Figure 4.**
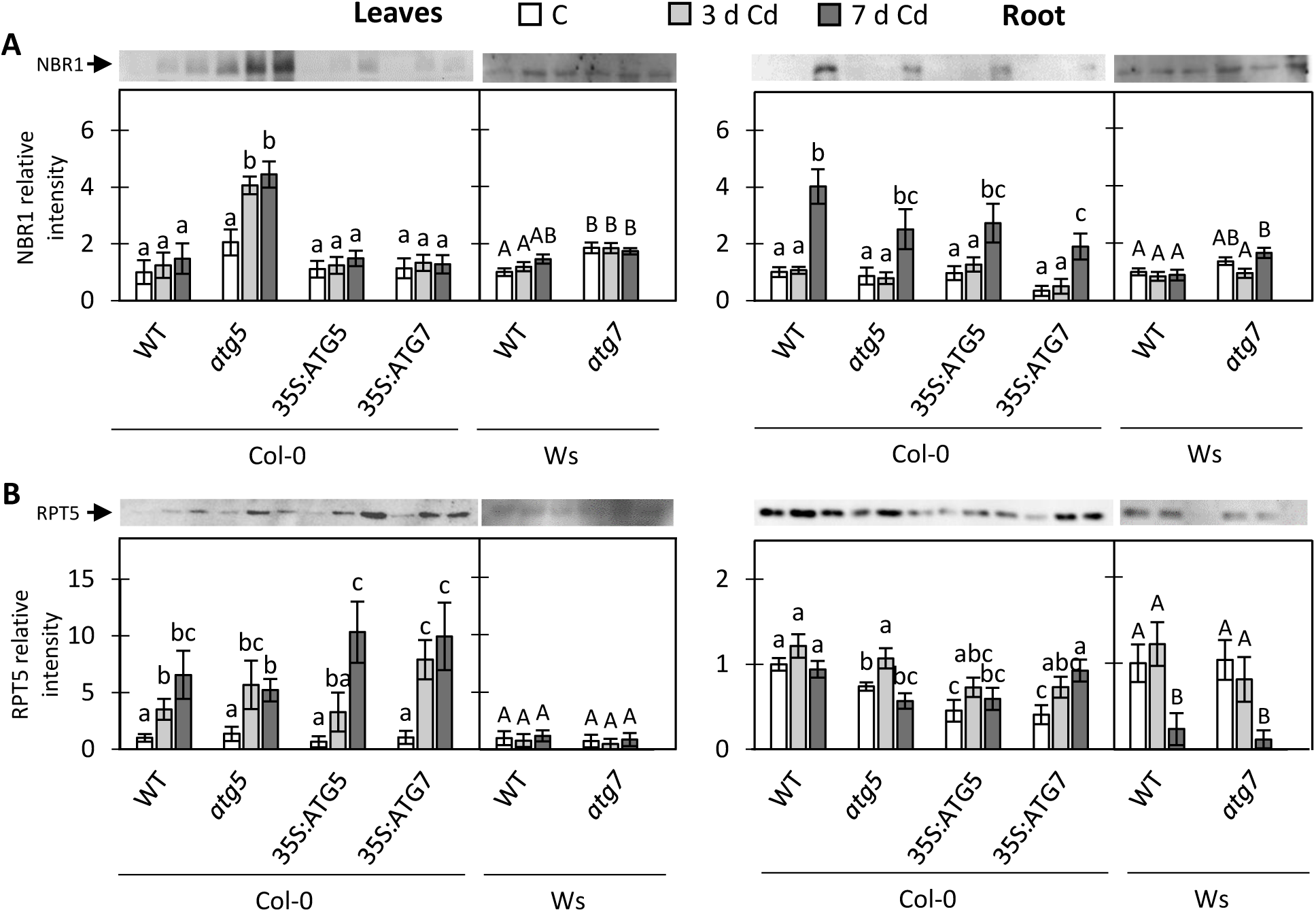
Cadmium effect on NBR1 and the proteasome unit RPT5 accumulation in leaves and roots after Cd treatment. Arabidopsis plants were treated with CdCl_2_ (50 µM) for 3 and 7 d and protein extract were obtained **A**, Western blot from proteins were incubated with anti-NBR1, 100 kDa. **B**, Western blot from proteins were incubated with anti-RPT5, 48 kDa. Histograms represents the band intensity mean ± SEM of 3 independent replicates. Values followed by different letters (lowercase letters compare Col-0 backgrounds, while capital letters compare Ws backgrounds) are statistically significant at p < 0.05.

Proteasome can act synergistically to autophagy to remove damaged proteins (Havé et al., 2018). The components of 26S proteasome, RPT5A subunit, was analysed by Western blot using specific antibodies. In leaves, RPT5A increased drastically in a time-dependent manner in all lines with Col-0 background but remained unchanged in lines with Ws background (Fig 4 B). In roots, no clear trends were observed for plants with Col-0 background, while RPT5A content was significantly diminished after 7dpt in both Ws WT and *atg7 roots* (Fig. 4B).

### Cd effect on phenotype and growth in wild type plants and autophagy-related mutants

We then analyzed the effect of Cd on growth and phenotype in WT and Arabidopsis mutants showing disturbances in autophagy. Exposure to Cd reduced root length in all genotypes, with strongest effect being in *atg5* mutants (Fig. 5A, B). The same was true for plant biomass. The critical role of *ATG5* was then confirmed in experiments with hydroponically grown plants (Fig. 6). Cd exposure resulted in a time-dependent decrease of plant growth and pigment content in all genotypes. The effect was less pronounced in plants with Ws background, and plants lacking functional *ATG5* were the most affected.

**Figure 5.**
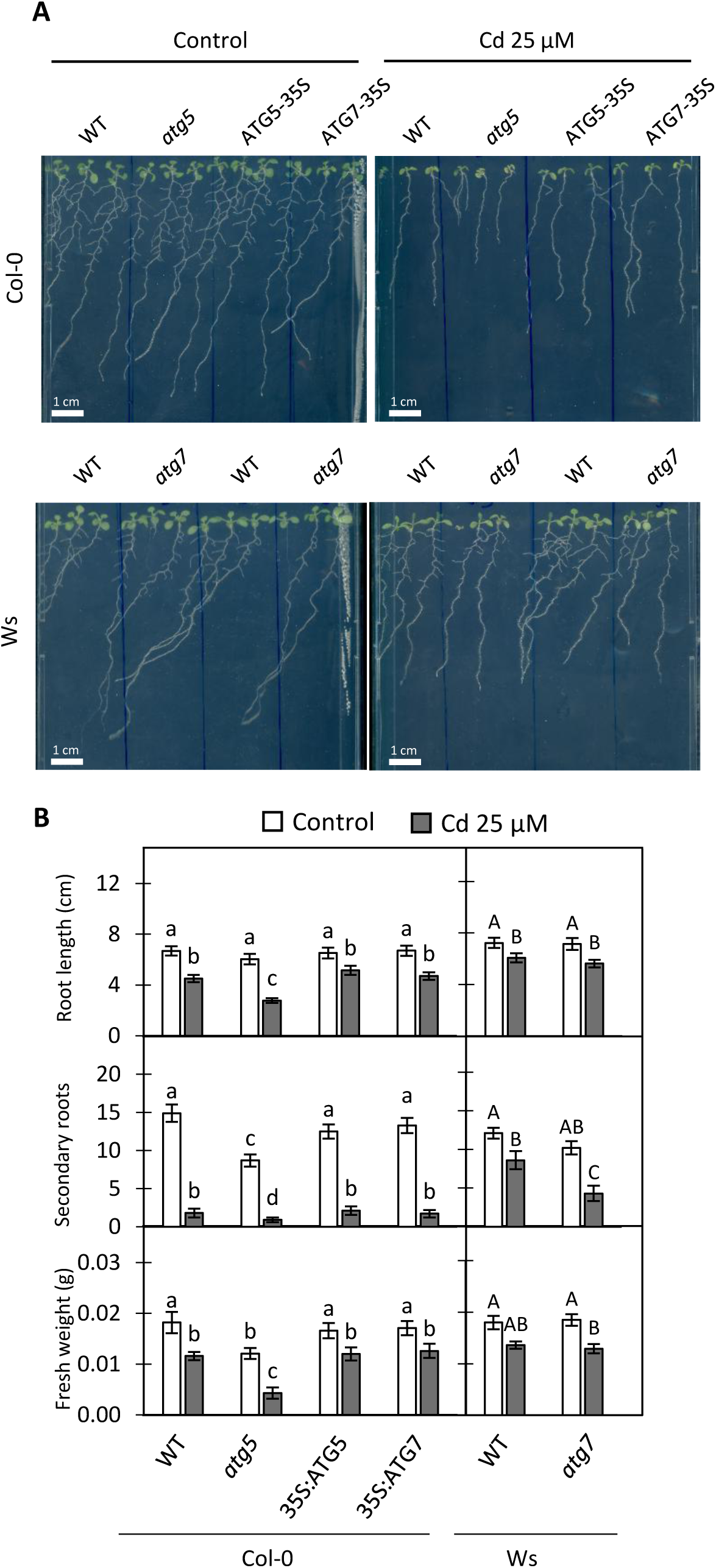
Cadmium effect on root WT plants and autophagy mutants. **A**, WT and *atg* mutants were grown for 12 d with CdCl_2_ (25 µM) and without Cd (Control). Image of root growth. **B**, Phenotype parameters: root length, secondary roots and fresh weight. Histograms represents mean ± SEM of 9 replicates. Values followed by different letters (lowercase letters compare Col-0 backgrounds, while capital letters compare Ws backgrounds) are statistically significant at p < 0.05.

**Figure 6.**
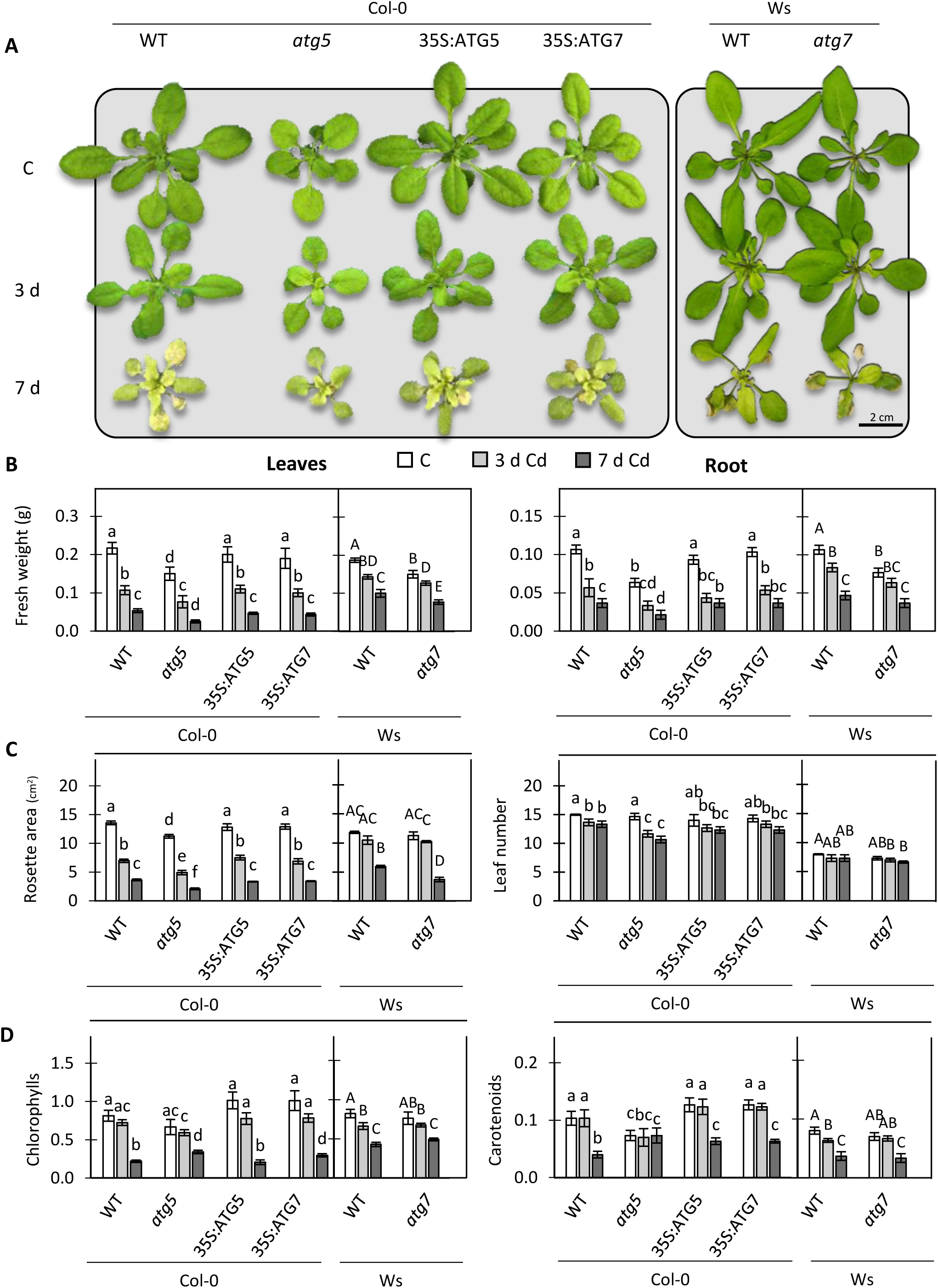
Cadmium effect on phenotype of WT and ATG-related mutants. **A**, Image of 20 days old plants grown in hydroponic containing Hoagland 0.5x medium with and without Cd (C). **B**, Fresh weight of leaves and roots. **C**, Rosette and leaf number. **D**, chlorophylls and carotenoid content. Histograms represents mean ± SEM of 6 replicates (three experiments with eight biological replicates each). Values followed by different letters (lowercase letters compare Col-0 backgrounds, while capital letters compare Ws backgrounds) are statistically significant at p < 0.05.

### Cadmium-dependent disturbances on ROS metabolism in WT and ATG-related mutants

One of the primary effects of Cd on plants and other organisms is the production of ROS and the induction of oxidative stress. Accordingly, we analyzed H_2_O_2_ and O_2_^.-^ production, oxidative parameters and antioxidant defenses in WT and ATG mutants in leaves. DAB staining revealed an increase in H_2_O_2_ content in leaves, with no significant difference between WT (Col-0) and mutant lines (Fig. 7A). Similar results were obtained by quantifying H_2_O_2_ content by biochemical methods (Fig.7 B). In roots the highest content of H2O2 was observed in *atg5*, meanwhile the WT (Col-0) and the over-expressors followed a similar pattern and Ws lines (WT and *atg7*) did not show changes in H_2_O_2_ content in response to Cd (Fig. 7 B). Superoxide content (measured by NBT staining) increased in leaves from Col-0 and *atg5* with the Cd treatment but was significantly lower in the over-expressors (Fig. 7 A). No Cd-induced increase in O ^.-^ was observed in lines with Ws background. The highest increase in MDA content (a lipid peroxidation marker) was in *atg5* line (Fig. 7 B).

**Figure 7.**
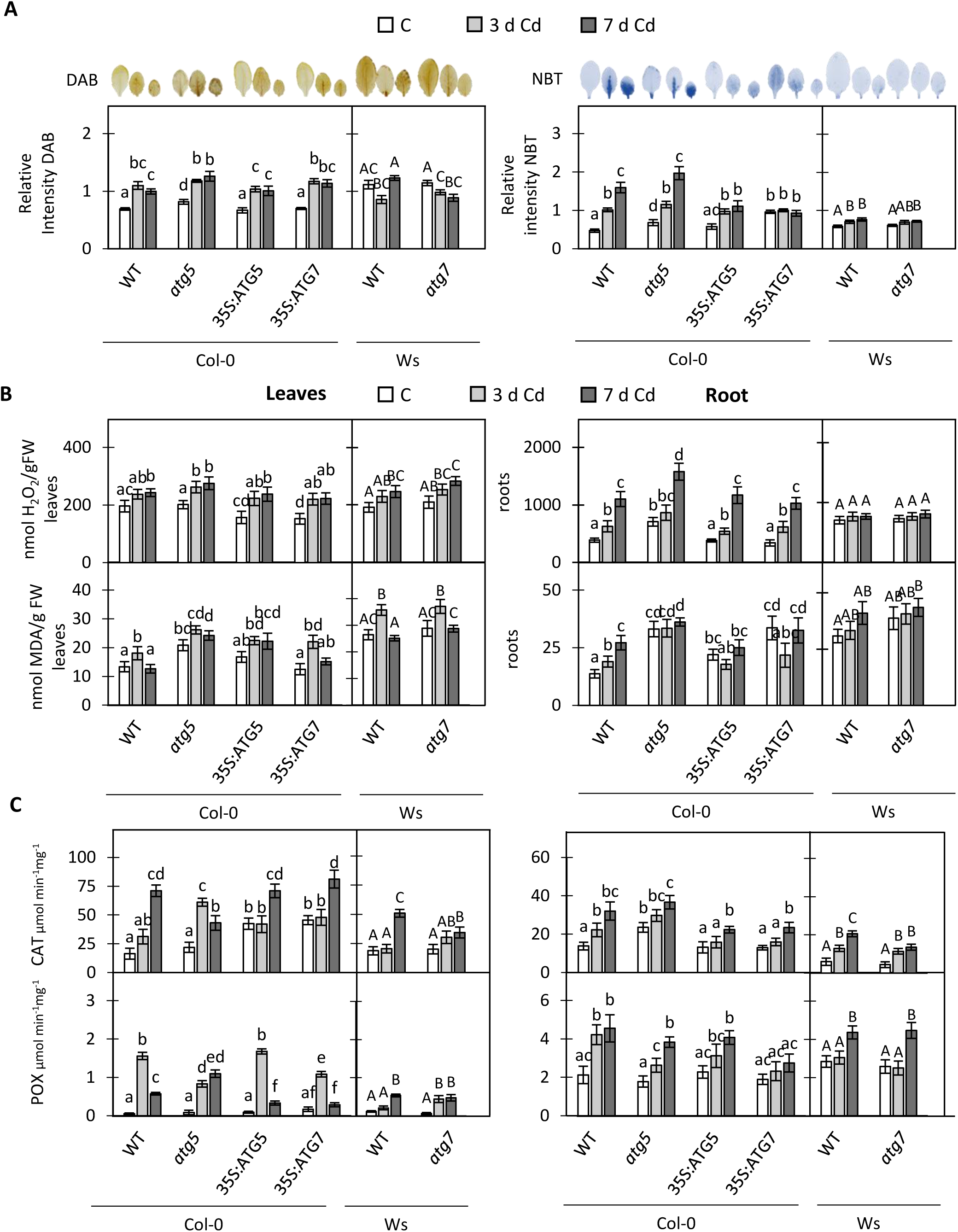
Cadmium effect on oxidative parameters and antioxidant defenses of WT and ATG-related mutants. Plants were grown in hydroponic conditions containing Hoagland 0.5x medium with (Cd, 50 µM; for 3d and 7d) and without Cd (C). **A**, Histochemical assays for H_2_O_2_ and O_2_^.-^ accumulation in leaves by using DAB and NBT staining, respectively. **B**, H_2_O_2_ concentration analysed by fluorimetry in leaves and roots. Lipid peroxidation analyzed as MDA content in leaves and roots. **C**, Enzimatic activities of catalase (CAT) and guayacol peroxidase (POX) in leaves and roots. Histograms represents mean ± SEM of 3 replicates. Values followed by different letters (lowercase letters compare Col-0 backgrounds, while capital letters compare Ws backgrounds) are statistically significant at p < 0.05.

Analysis of CAT activity did not reveal any significant differences between genotypes neither in leaves nor roots, with time-dependent increase in enyme activity in all lines (Fig. 7 C). Guaiacol peroxidase activity (POX) increased transiently by day 3 and then reduced in Col-0 leaves but stayed elevated in roots (Fig. 7 C).

### Role of autophagy in metal and ion homeostasis under cadmium stress *Ionomic profile*

To determine if autophagy regulates the uptake of Cd and other metals and ions, we analysed the elemental composition of *atg5* and *atg7* mutants with their respective WT after 3 days of treatment, when the induction of autophagy was apparent in both leaves and roots. A 3-d exposure to Cd resulted in a significant (⁓ 1.5-1.6 folds) decrease in the content of all essential macronutrients (K, P, Ca, Mg, S) in leaves of all lines (Table 1). In roots, a decline in K content was even more pronounced (⁓ 2 folds) but the content of other macronutrients even increased slightly, suggesting effects of Cd on xylem nutrient loading and their delivery to the shoot. Also decreased was the content of essential micronutrients (Cu, Fe, Mn, Zn) in leaves while in roots it was either unchanged or slightly increased. No clear difference in ionome profiles between wild types and respected autophagy lines was observed (Table 1) except for Cd which was always highest in roots of WT lines. *Agt5* mutants had highest Cd content in the shoot, of all genotypes studied. Plants with Ws background accumulated higher content of Cd in roots, that this ecotype has developed efficient mechanisms to accumulate Cd (Amaral dos Reis et al., 2021; Park and Ahn, 2017).

**Table 1.**
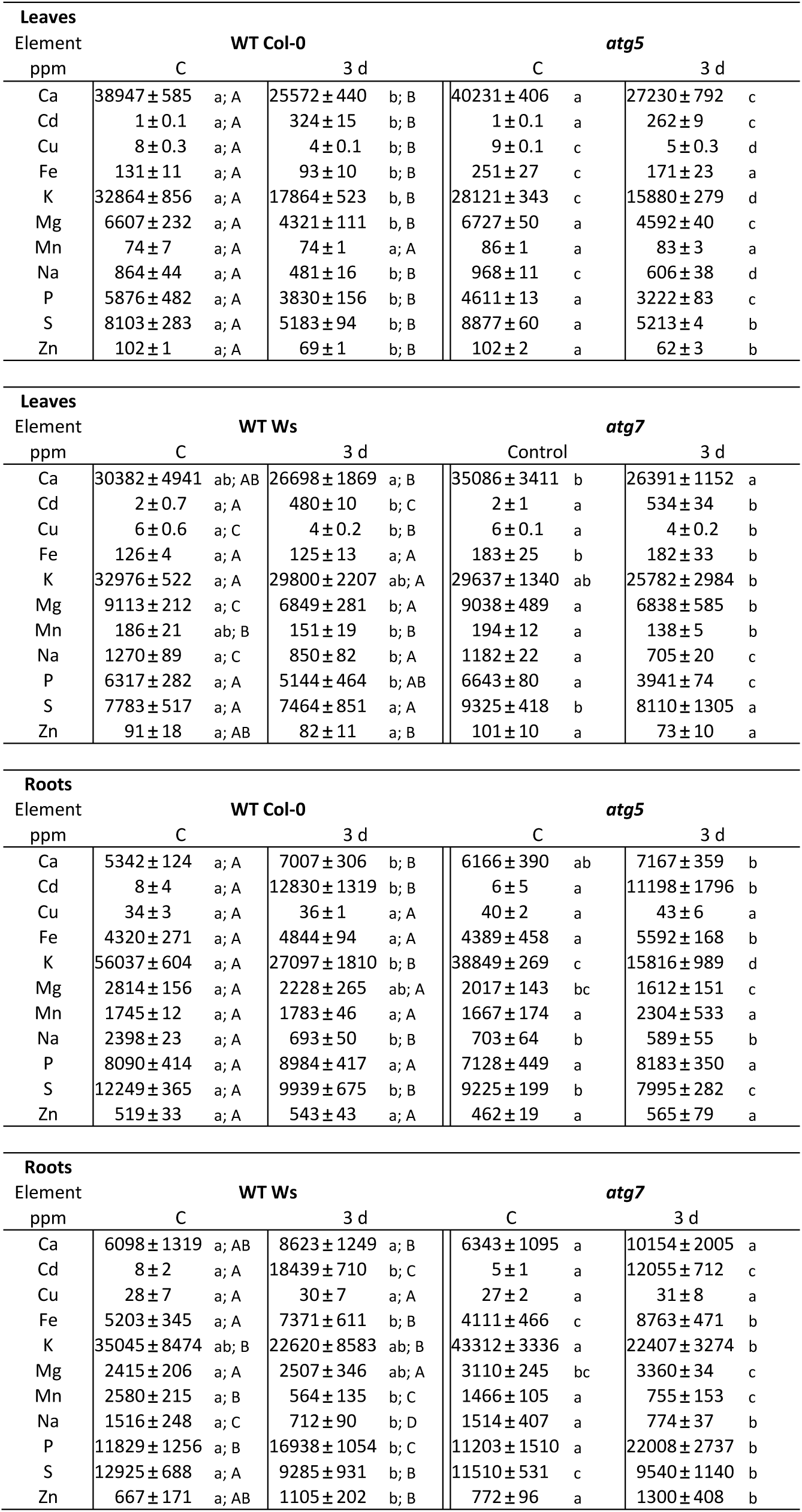
Cadmium and autophagy disturbances effect on ionomic of leaves and roots of plant untreated or treated with cadmium during 3 days. The analyses were carried out by inductively coupled plasma mass spectrometry (ICP-MS). Values followed by different letters (lowercase letters compare same backgrounds lines, while capital letters compare Col-0 and Ws backgrounds) are statistically significant at p<0.05

Suppl. Fig. S2 shows a heat map depicting the distribution of metals and other elements analysed in the different genotypes and treatment. The results showed a good separation of leaves and roots and control and treated plants. The highest content of S, K^+^, Cu^2+^ and Mn^2+^ was associated with control plants independently of the genotype, while the highest content of Cu^2+^, Cd^2+^ and Fe^3+^ was found in Cd-treated roots. The highest correlations were observed between *atg7* and its WT in roots for Cd^2+^, P, Fe^3+^ and Zn^2+^.

#### Net Cd uptake by roots

We then analysed the flux of Cd^2+^ in the elongation root zone of Arabisopsis plants. Net Cd^2+^ uptake was stronger in *atg5* mutants compared with its WT (Col-0) while no such difference was found in *atg7* line in Ws background (Fig. 8). Cd^2+^ uptake was prolonged in *atg5* and still significantly higher than in WT after ⁓20 min of exposure (Fig 8B).

**Figure 8.**
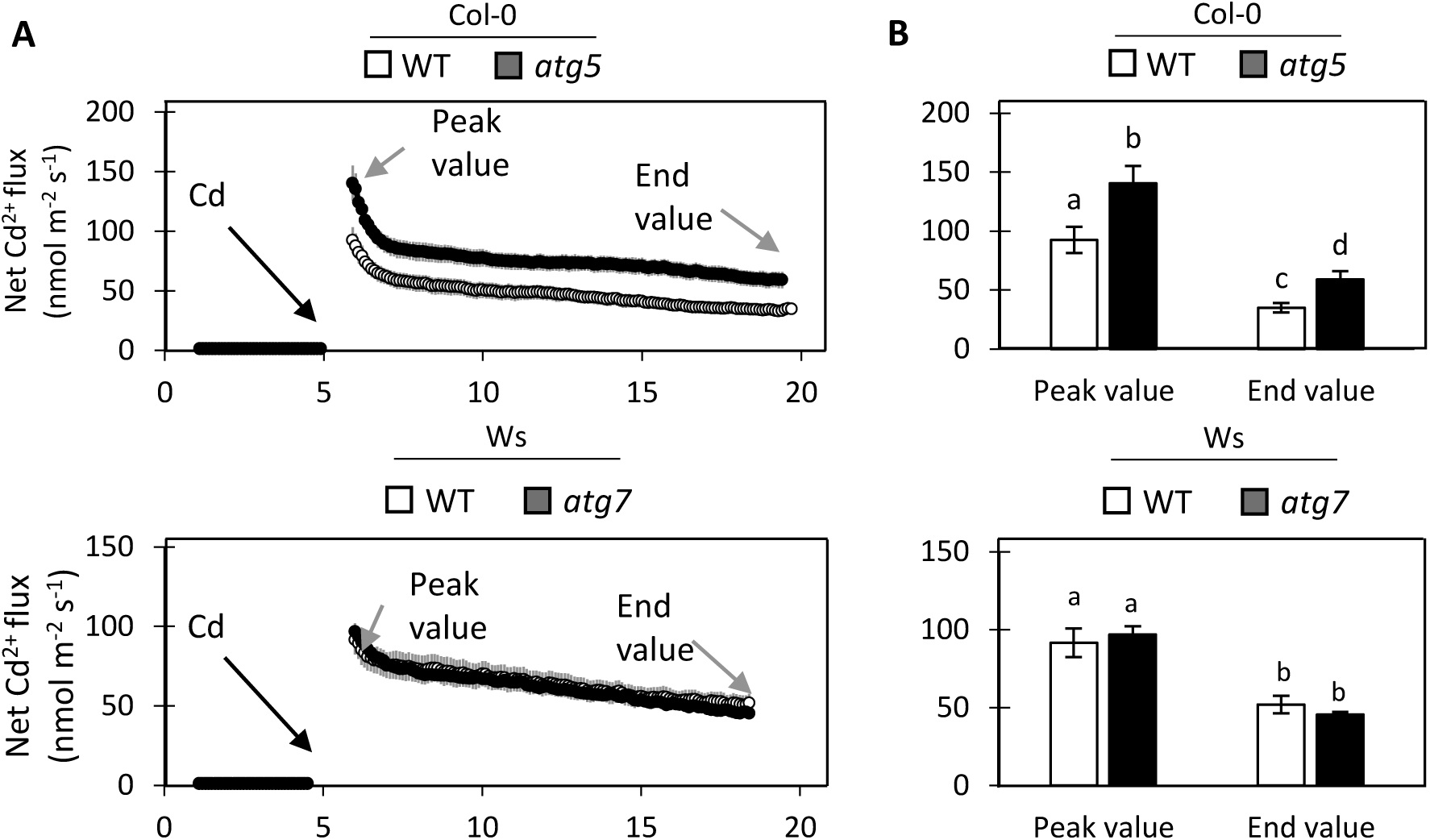
Cadmium Flux changes in autophagy mutants. **A** Net Cd^2+^ fluxes measured from epidermal root cells in the elongation zone of Col-0: WT and *atg5*, and Ws: WT and *atg7* plants in response to 50 M µCd^2+^ added after 5 minutes (black arrow). **B** Comparative analysis of the magnitude of net Cd^2+^ flux responses (peak and end). Values followed by different letters (lowercase letters compare Col-0 backgrounds, while capital letters compare Ws backgrounds) are statistically significant at p < 0.05.

#### Transcriptomic analysis of metals and K transporters

To understand the changes observed in ionomic data, we have analysed the expression of some metal transporters involved in their uptake and translocation. In *leaves HMA2* and *HMA4* Fe^3+^ transporters (which can also transport Cd) were significantly up-regulated in a time-dependent manner (Fig 9); effects were less pronounced in WT lines and much stronger in mutant lines. Expression of another Fe^3+^ transporter, *IRT3* (which can also transport Cd) peaked at day 3 and declined afterwords. No clear difference between ecotypes was observed. *YSL3* gene which mediate long-distance Cd transport (Tao and Lu, 2022) was also significantly upregulated, peaking at 3 d in lines with Col-0 background and 7 d for Ws background. Again, effect was less pronounced in WT lines compared with appropriate mutants (Fig 9). Cd exposure also transiently upregulated expression of *ZIP5* (involved in Fe^3+^, Zn^2+^ and Cd uptake in the roots (Lin et al., 2016; Tan et al., 2020), with strongest effect observed in *atg5* mutant. *NRAMP3* and *NRAMP6* genes that encode several metals and Cd efflux transporters on tonoplast and ER, respectively, were upregulated in a time-dependent manner in all lines with Col-0 background. In Ws background, no clear trends were observed.

**Figure 9.**
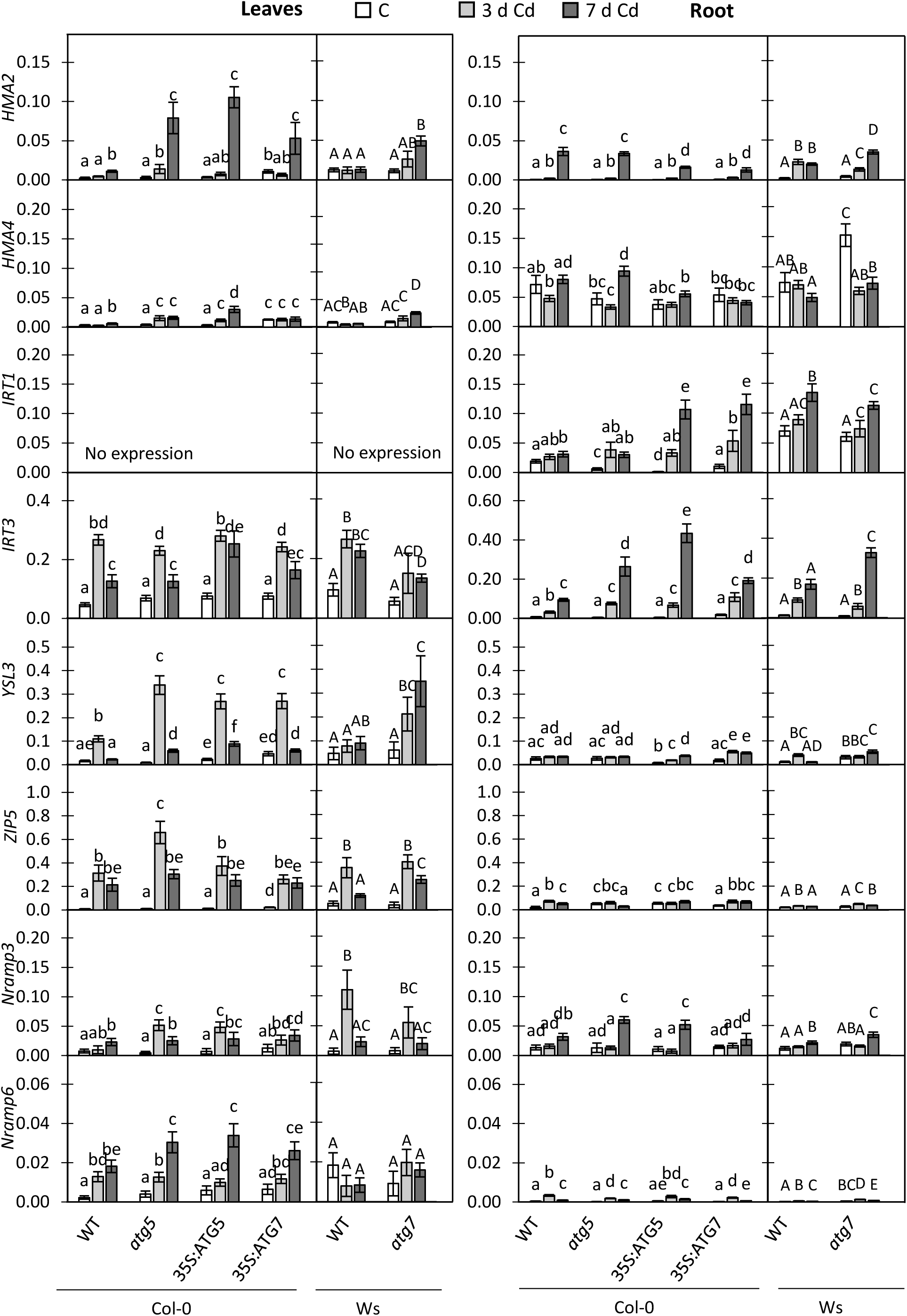
Cadmium effect and autophagy regulation of metal transporters genes expresión in leaves and roots. Plants were grown in hydroponic conditions containing Hoagland 0.5x medium with (Cd, 50 µM; for 3d and 7d) and without Cd (C) and metal transporter expression in roots and leaves was analysed. Histograms represent mean ± SEM of 3 replicates gene expression compared with *TUB4*. Values followed by different letters (lowercase letters compare Col-0 backgrounds, while capital letters compare Ws backgrounds) are statistically significant at p < 0.05.

In *roots*, *HMA2* was significantly upregulated in all lines, with highest expression being at day 7 (Fig. 9,) while *HMA4* expression was largely unaffected. *IRT1* transcripts (specific for roots only) were also highest at day 7, and so were those for *IRT3*, with the highest levels observed in over-expressor lines (Fig 9). Changes in transcript levels of *YSL3* were minor and physiologically insignificant. *NRAMP3* was upregulated in time-dependent manner in all lines but transcript levels of *NRAMP6* were mostly unaffected.

Given a causal link between cytosolic K^+^ content and cell apoptosis (Shabala, 2017), we then looked at expression of HAK5 (mediating high-affinity K^+^ uptake by roots) (Ródenas et al., 2021) and GORK (encoding outward-rectifying K^+^-efflux channel) (Adem et al., 2020). As expected, Cd-induced changes in *HAK5* expression in leaves were physiologically non-essential (Fig 10), while in roots Cd treatment has caused a very major upregulation of *HAK5* transcripts, peaking at day 3, with strongest effects observed in WT plants (of both backgrounds). *GORK* transcripts were massively upregulated (by 5 to 10-folds) in both leaves and roots, with no clear difference between genotypes. A heat map showing relative transcript levels of differentially expressed ion/metal transporter genes across genotypes is presented in Suppl. Fig. S4.

**Figure 10.**
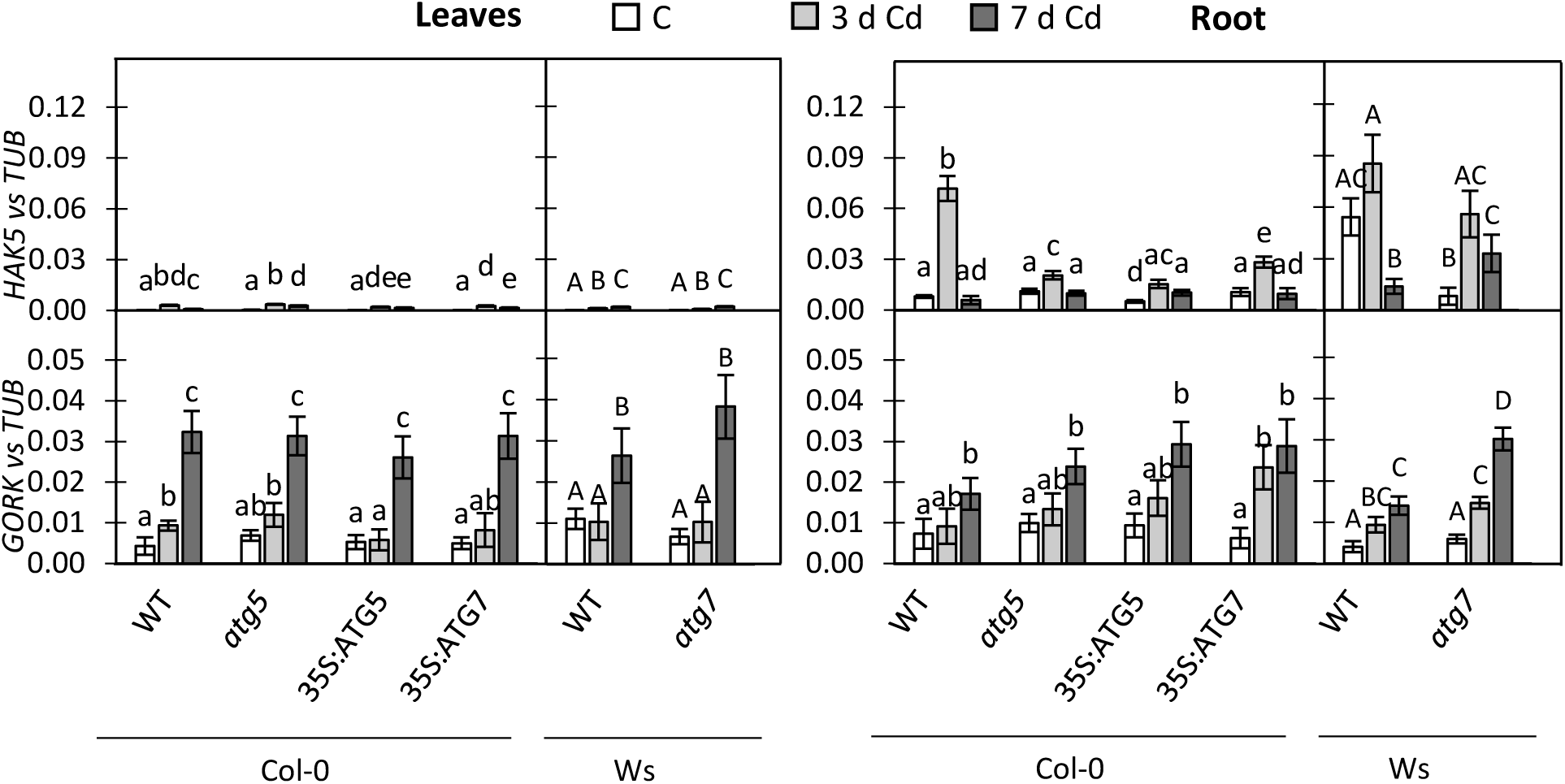
Cadmium effect and autophagy regulation of potassium transporters genes expresión in leaves and roots. Plants were grown in hydroponic conditions containing Hoagland 0.5x medium with (Cd, 50 µM; for 3d and 7d) and without Cd (C) and potassium transporter expression in roots and leaves was analysed. Histograms represent mean ± SEM of 3 replicates gene expression compared with *TUB4*. Values followed by different letters (lowercase letters compare Col-0 backgrounds, while capital letters compare Ws backgrounds) are statistically significant at p < 0.05.

## Discussion

Autophagy is a well know process involved in recycling cell components under nutrient starvation, to sustain plant growth and development (reviewed in Petersen et al., 2024 (Petersen et al., 2024a)). However, the role of autophagy in plant responses to an excess of heavy metals and, specifically, to Cd has been less explored. In human and mammals heavy metals can modulate autophagic flux by either inducing this process as a protective response, blocking it, or even switching its protective role toward a pro-cell death mechanism (Genchi et al., 2020; Martínez-García and Mariño, 2020; Rahman et al., 2023). This study showed that acute Cd treatment induced autophagy in a time-and concentration-dependent manner in both leaves and roots of Arabidopsis plants. As expected, this process occurs first in the roots, considering that roots are more directly exposed to Cd. Both roots and shoots shown a presence of autophagosomes in response to Cd, although their density were considerably smaller than those observed under nutrient deprivation such as P (Naumann et al., 2019) or reduced Carbon sources (Kacprzak and Van Aken, 2022; Laureano-Marín et al., 2016). Apple plants exposed to Cd also show an increase of vesicles in the vacuole identified by the authors as autophagosomes by electron microscopy (Huo et al., 2022), although no confirmation by fluorescence associated to AGT8 or immunogold analysis and autophagy flux analysis have been carried out in this study (Huo et al., 2022).

The transcriptomic analyses of autophagy markers such as *ATG4a, ATG4b, ATG5, ATG7, ATG8a* and *ATG8h* in roots and leaves shown here reinforce the Cd-dependent activation of autophagy, although differences in time-course regulation between some of these markers were observed in leaves and roots. Interestingly, both Ws background plants (WT and *atg7*) did not significantly change the expression of most ATGs and *TOR* analyzed in either leaves or roots in response to Cd but had constitutively higher expression levels of most ATGs analysed under control conditions. This result suggests that the Ws background has probably already induced the autophagy process which could explain the higher tolerance of this genotype against metals. Additionally, the reduced effect of Cd on *ATGs* expression in Ws could be related to the different tolerance of this ecotype to metals being more tolerant to Cu and Cd than Col-0 genotype. The tolerance to Cu in Ws was associated with the upregulation of the Cu-chelating metallothionein 2 (MT2) (Murphy and Taiz, 1995) and the downregulation of Cu transporter (COPT) genes (Amaral dos Reis et al., 2021), as well as differential potassium flux (Murphy et al., 1997). In the present study Ws ecotype accumulated more Cd in roots than Col-0 but, as reported previously (Amaral dos Reis et al., 2021), this ecotype was more efficient in sequestering the metal to prevent its toxicity. This could explain the absence of changes in ATGs expression in this genotype in response to Cd. Additionally, differential gene expression of *HMA* transporter genes in both ecotypes could explain the higher sensitivity of Col-0 to metals (Park et al., 2012). Amaral dos Reis et al (2018) (Amaral dos Reis et al., 2018) have also reported that leaves of both accessions use different strategies to cope with metal stress, with Ws plants investing mainly in maintaining nutrient homeostasis, while Col-0 plants are more focused on metal detoxification strategies.

TOR has been established as having a central role for integrating external and internal metabolic cues and there is much evidence to support its role in regulating autophagy (Burkart and Brandizzi, 2021; Dong et al., 2022; Mugume et al., 2020). In the present study, TOR was up-regulated in all Col-0 genotypes, mainly in both over-expressor lines, which could be a mechanism to regulate excess of autophagy, while minor changes were observed in Ws WT and *atg7*. The changes observed in TOR expression in both genotypes, Col-0 and Ws in the present work correlated with the differential changes observed in the ion homeostasis in both genotypes. This result could be explained by the TOR signalling participation in nutrient sensing and deficiency (Fu et al., 2020). However, according to Pu and Soto-Burgos (Pu et al., 2017; Soto-Burgos and Bassham, 2017) TOR would not be involved in oxidative stress and ER stress triggering autophagy processes. In *Schizosaccharomyces pombe* yeast it has been reported that under Cd toxicity downregulation of TOR1 improved growth rate conditions and antioxidant defence mechanisms and the Cd uptake was markedly smaller than in wild-type cells, although these authors did not establish any relationship with autophagy (Požgajová et al., 2020). These authors also showed that TOR1 downregulation promoted the accumulation of metals such as Cu and Fe (Požgajová et al., 2020).

We have also carried out a Western blot analysis to look for changes in ATG4, ATG8 and ATG5 accumulation during the Cd treatment in WTs (Col-0 and Ws) and the different *atg* mutants analysed. ATG8 accumulation in WT (Col-0) under Cd treatment matches with the upregulation of *ATG8a* and the activation of autophagy reported by the increase of lipidated ATG8a observed by Western blot. Interestingly, in *atg5* a strong accumulation of ATG8 was observed, even under control conditions, as a result of the inhibition of autophagy because ATG5 is necessary for the activity of the ATG12 complex. A higher abundance of ATG8, showing different bands, including lipidated ATG8 were also observed in the *atg5* and *atg7* in response to extended darkness as the result of *ATG8* up-regulation (Thompson et al., 2005), although in our study no significant up-regulation of *ATG8a* and a slight decrease of *ATG8h* transcript was observed in *atg5*. However, in *atg7* no significant changes were observed in terms of transcription but a significant decrease of ATG8 accumulation was observed. Arabidopsis *atg2* mutants showed an accumulation of ATG5, ATG8A, ATG18A and ATG12 in response to high temperature and light (Jiang et al., 2020).

NBR1 is a plant aggrephagy receptor essential for maintaining proteostasis under both heat stress and non-stress conditions (Jung et al., 2020) and it could also participate in the response to Cd treatment to judge by the slight increase, mainly in roots, observed in the present work in response to Cd. Jung et al (2020) (Jung et al., 2020) have reported that NBR1 is required for the heat-induced formation of autophagic vesicles. Interestingly, the content of NBR1 considerably increased in *atg5* mutants, as a result of blocking autophagy, and similar results were obtained in *atg7* seedlings in response to heat stress (Jung et al., 2020). In turn, Cd did not significantly affect the NBR1 content neither in roots nor in leaves from WT Ws background and a small but significant increase of NBR1 took place in *atg7* line. Meanwhile, overexpression of *ATG5* and *ATG7* showed similar accumulation than in its WT in Col-0 background. These results support that NBR1 and its substrates could be degraded by autophagy as suggested by Svenning et al (2011) (Svenning et al., 2011) and Jung et al (2020) (Jung et al., 2020), although this requires deeper research. Currently, the functions of the plant NBR1 under heavy metal stress remain elusive, although it has been suggested that it could participate in pexophagy regulating the population and quality of peroxisomes in response to Cd stress (Calero-Muñoz et al., 2019). Proteasome also appear to be involved in cell response to Cd, as seen by the accumulation of RPT5A in leaves. However, RPT5A accumulation was not apparently regulated by autophagy in leaves, as indicated by the lack of changes in response to Cd in the *atg5* line, although it was significantly increased in the overexpressors. Interestingly, *atg7* and its WT (in Ws background) kept low levels of RPT5A in leaves and followed an opposite pattern in roots with a decrease at 7dpt, which could be due to the lower effect of Cd in these genotypes. Our results therefore suggest that there are no compensatory changes in proteasome under autophagy deficiency. However, this compensatory effect was observed in Arabidopsis mutants *atg5* during ageing by Havé et al (2018) (Havé et al., 2018). Moreover, the ubiquitin-proteasome system (UPS) was reported as a putative interactor of ATG5 (Elander et al., 2023). Additionally, overexpression of *Ubiquitine2* from tobacco in Arabidopsis increased Cd tolerance by decreasing Cd accumulation, as a result of the higher expression of the Cd/metal exporter *PDR8*, and the lower expression of the xylem-loading exporters *HMA2* and *HMA4* (Bahmani et al., 2019). Additionally, A lower expression of the importer *IRT1* and a higher 20S proteasome activity was also observed (Bahmani et al., 2019). However, the specific mechanism has not been stablished so far, and further analysis is required to establish the relationship between proteasome and autophagy in response to Cd.

### Plant growth, oxidative stress and autophagy

Plant growth analysis demonstrate that disturbance in autophagy (*atg5*) negatively affected the growth mainly under Cd stress and that induction of autophagy in over-expressors did not significantly improve the tolerance to this heavy metal (Fig. 5 and 6). Previous studies suggested that nutrient-starved plant species show more sensitive phenotype in autophagy-disrupted mutants such as *atg5* (Zou et al., 2025) or *atg6* (Cao et al., 2022), while overexpression of *ATG5* or *ATG7* stimulates ATG8 lipidation, autophagosome formation, and autophagic flux, favouring stress resistance and the growth in Arabidopsis plants (Minina et al., 2018).

In contrast to our data, Huo et al., (2022) (Huo et al., 2022) reported that overexpressing ATG10 in *Malus* plants reverted the Cd-dependent reduction of growth, presumably due to the reduction of Cd uptake and translocation, and the improvement of antioxidative defences (Huo et al., 2022). However, none of these changes have been observed in the present study. Notably, being exposed to a much higher Cd concentration (120 μM vs 50 μM in our work), *Malus* accumulated around 22- and 13-time less Cd in stem and roots, respectively. Therefore, under our experimental conditions the induction of autophagy appears not to be a protective mechanism against Cd stress but its consequence. We can speculate that Cd exposure can give rise a differential response depending on the plant species and the metal concentration. Interestingly, Ws lines showed a higher fresh weight, rosette area and greener with higher chlorophyll content, and lower content of carotenoids than Col-0 genotypes. In microalgae *Crypthecodinium* sp and *Chlamydomonas* cells, a link has been established between photo-oxidative damage, ROS accumulation, autophagy activation and the reduction of carotenoid content (Li et al., 2024; Pérez-Pérez et al., 2012; Tran et al., 2019). However, the relationship between carotenoids and autophagy has not been explored in Arabidopsis plants.

Much evidence supports the theory that oxidative stress can trigger autophagy in response to abiotic stress such as drought (Liu et al., 2009; Tang and Bassham, 2022) and salinity (Luo et al., 2017) and autophagy is required for tolerance to both abiotic stress (Liu et al., 2009; Luo et al., 2017). Therefore, oxidative stress can regulate autophagy, but autophagy could also regulate the cell response to oxidative stress. In this context, the overexpressing *thioredoxin o1* in tobacco BY-2 cells (PsTRXo1) showed a higher gene expression, protein content and activity of ATG4, as well as an increase of protein content of ATG8, and lipidated ATG8 than WT cells, supporting the involvement of autophagy in the response of the mutant against H_2_O_2_ (De Brasi-Velasco et al., 2021). Additionally, Arabidopsis *atg2* and *atg7* mutants contained more oxidized proteins and were hypersensitive to both salt and osmotic stresses (Luo et al., 2017) and RNAi-ATG18a Arabidopsis mutants were more sensitive to oxidative stress (Xiong et al., 2007). In the present study, Cd induced oxidative stress in all Col-0 genotypes, although over-expressors showed slightly lesser accumulation of H_2_O_2_ and O_2.-_ in leaves, thus suggesting that induction of autophagy is not enough to protect plants from oxidative damage derived from acute Cd treatment exposure. Interestingly, Ws lines were apparently more resistant to Cd because they did not show oxidative stress by the metal neither in the leaves nor in the roots. This process was not conferred by higher activity of enzymatic antioxidants, as shown by analysis of CAT and POX activity (Fig 7). Arabidopsis *atg12* mutants didn’t show significant changes in some of the main antioxidants involved in Glutathione-Ascorbate cycle and catalase during the maturation process, which is linked to oxidative stress, and even the content of H_2_O_2_ was reduced in *atg12* mutants (Barros et al., 2023). Therefore, a direct link between antioxidant defences, ROS production and autophagy is not always clear. However, in *Malus* plants the overexpression of *ATG10* protected plants from oxidative stress induced by Cd by improving antioxidants and decreasing O_2.-_ and H_2_O_2_ accumulation (Huo et al., 2022). Differences between both species could be related to different strategies associated to the accumulation of Cd in a nontoxic way and other factors to be stabilised. Additionally, increasing evidence supports the redox control of autophagy, although the molecular mechanisms underlying this regulation have not been fully elucidated in plants. Thus ATG4 has been shown to be a target of H_2_O_2_-dependent posttranslational modifications in humans (Scherz-Shouval et al., 2007), yeast (Perez-Perez et al., 2014), algae (Pérez-Pérez et al., 2016) and plants with oxidation of ATG4 giving rise to the inactive form (Pérez-Pérez et al., 2021). Recently, ATG3 activity from yeast and algae has been demonstrated to be also regulated by redox changes (Mallén-Ponce and Pérez-Pérez, 2023). Therefore, in the present study, the absence of improvement response in over-expressors could be related to overoxidation of ATG4 or other autophagy targets thus blocking autophagy under certain conditions. This suggestion could be supported, at least in roots, by the accumulation of ATG8 free and ATG8-PE at 3 and 7 dpt.

### Cd and ion homeostasis regulation by autophagy

Cadmium treatment-induced disturbances in ion homeostasis in both roots and shoots were mainly associated with a decrease in the content of most of the ions analysed. Cd may cause disturbances in the homeostasis of essential ions by competition with the transporters (Gupta et al., 2017; Haider et al., 2021). An example is the Cd-induced deficiency of Zn and Cu levels in shoots and roots observed in Arabidopsis plants (Gielen et al., 2017; Gupta et al., 2017) and the Cd-induced deficiency of Fe (McInturf et al., 2022). Much evidence suggests that these changes are regulated by ROS induced by Cd (Hafsi et al., 2022; McInturf et al., 2022; Sandalio et al., 2023). Disturbances in autophagy in *atg5* significantly alters ion accumulation in the leaves and roots of Arabidopsis Col-0. Thus, *atg5* showed a slight decrease of Cd^2+^ content in leaves but did not affect its content in roots. Additionally, Cd increased Fe^3+^ and decreased K^+^ and P content in leaves from *atg5*, while it increased Fe^3+^ and reduced K^+^, Mg^2+^ and S in roots. Meanwhile, Ws lines accumulated a higher amount of Cd and most of the ions, in comparison with Col-0. These results suggest that autophagy can regulate ion homeostasis in plants. In fact, in *Malus* plant overexpressing *ATG10* a decrease of Cd content in shoots was observed (Huo et al., 2022). Shinozaki and Yoshimoto (2021) (Shinozaki and Yoshimoto, 2021b) have reported the induction of autophagy in response to excess of Zn^2+^ and *atg5* were more sensitive to Zn^2+^ excess but this negative effect could be alleviated by supplementation with high levels of iron to the media (Shinozaki et al., 2021). These authors also reported an excess of Zn^2+^ in *atg2* and *atg5* mutants that led to chlorophyll degradation and chlorosis (Shinozaki et al., 2021). Meanwhile, in our study Zn^2+^ content did not show significant changes in *atg5,* neither under control conditions, nor under Cd stress in leaves and roots.

In mammal cells it has been suggested that ion homeostasis can regulate autophagy, and then autophagy can regulate different ion channels and transporters (Zhang et al., 2022). However, in plants this relationship has still not been clearly deciphered. In the present study all metal transporters analysed were up-regulated at 3 or 7 dpt in Col-0 plants and the pattern of expression differed in *atg* mutants independent of silencing and over-expression of *ATG5* and *ATG7*, which suggests that the activity of transporters could probably be regulated at different post-translational levels, most likely by ROS ( Sandalio et al., 2023).

Additionally, it was proposed that environmental stresses giving rise to loss of cytosolic K^+^ can induce autophagy and PCD in roots (Demidchik et al., 2017). This hypothesis is based on the results obtained in *gork1-1* under salinity in which a diminished K^+^ efflux was associated to fewer autophagosome formation in comparison to WT plants (Demidchik et al., 2010). Arabidopsis *kup8-2* plants, which suppressed the K^+^ transporter *KUP8* were reported to be more resistant against a mixture of heavy metals than WT plants (Sanz-Fernández et al., 2021), which could support an important role of K^+^ under abiotic stress conditions. However, in the present study Cd promoted a significant decrease of K^+^ in WT, which could therefore trigger the induction of autophagy. However, no significant changes were observed in *HAK5* and *GORK* expression between WT and *atg* mutants, therefore raising doubts about the role of K^+^ in autophagy induction. However, we cannot rule out changes in transporter activities due to ROS-dependent posttranslational modification as reported previously (Calero-Muñoz et al., 2019; Demidchik et al., 2017; Sandalio et al., 2023) or the involvement of other K^+^ transporters. The degradation of ion channels and transporters mediated by autophagy could be another possibility which deserves deeper analyses. In mammal tissues transporters and ion channel degradation by autophagy could be a feedback mechanism to control the excessive activation of autophagy during stress (Zhang et al., 2022). An example is the IRT1 which can act as a transporter and a receptor, being able to sense the excess of non-iron metal substrates in the cytoplasm, by regulating its own degradation (Dubeaux et al., 2018). Thus, the metal binding to a Hs-rich stretch of IRT1 promotes phosphorylation by CIPK23 kinase which in turn facilitates the recruitment of the IDF1 E3 ligase and the polyubiquitination which promote endosomal sorting and vacuolar degradation of IRT1 (Dubeaux et al., 2018). This mechanism allows iron uptake to optimize and prevent the accumulation of toxic metals in plants. Additionally, Chiu et al (2023) (Chiu et al., 2023) have reported that the degradation of Pi transporter PHT1;1/2/3 is enhanced in *atg5-1* under Pi deficiency.

Overall, it appears that the regulation of metal/ion accumulation by autophagy is a very complex process which will depend on the transcriptional regulation of ion/metal transporters but also on the post-translational regulation of these channels/transporters by ROS, NO and phosphorylation, among other modifications as well as the degradation of these transporters.

## Conclusions

High concentrations of Cd promote autophagy, as demonstrated by the increase of autophagy flux using different approaches as well as the upregulation of different ATGs. This induction could be regulated by oxidative stress and be a strategy to improve the adaptation of plants to Cd toxicity. Interestingly, genotypes with Ws background differ considerably to Col-0 being more resistant to cadmium. This is partially due to constitutive upregulation of *ATG4a*, *ATG4b*, *ATG5* and *TOR* and, hence, a higher level of basal autophagy. However, *atg7* plants followed a similar pattern of Ws in most parameters analysed, suggesting that autophagy can contribute but is not the unique determinant of the tolerance to high Cd concentrations in Ws. Additionally, our results suggest that NBR1 is required to keep proteostasis in response to Cd because suppressing autophagy could promote an increase of NBR1.In its turn, proteasome could collaborate with autophagy eliminating damaged proteins, although a clear collaboration of both processes in response to Cd is not clear, despite of changes observed in RPT5 content in *ATG5* and *ATG7* overexpressors. Further studies are required to clarify this issue Ionomic, flux analysis and transcriptional analysis of different transporters suggest that autophagy can regulate the accumulation of different metals both at transcriptional and posttranslational levels. The possibility of transporter degradation such as has been reported for IRT1, by proteasome or by autophagy cannot be ruled out and requires further analysis. Finally, although autophagy could protect plants against Cd, under acute Cd exposure the constitutive induction of autophagy fails in improving plant response to Cd in Arabidopsis plants.

## Authors contribution

LMS conceived the research, got the financial support and wrote the manuscript. AMCA, performed most experimental research. FLPG, JE, RMD, MCRP, SS and LMS contributed to some experiments. MCRP, SS and LMS discussed and edited the article. All authors contributed to the final version of the manuscript.

## Data availability

Data will be made available on request and through the CSIC repository

## Acknowledgements

AMCA, MCRP and LMS were supported by grants P20_00364 from the Junta de Andalucía, and PID2021-122280NB-I00 from the Ministry of Science, and Innovation and Universities, the ‘Agencia Estatal de Investigación’ and the European Regional Development Fund co-funding (MCIU/AEI/ERDF). JE was supported by a fellowship for academic staff from the Junta de Andalucía (PREDOC_00917). Authors acknowledge Dr. David Porcel from the Microscopy Confocal facilities from the University of Granada, where confocal studies were carried out.

## Abbreviations

ATG: autophagy related genes;
CAT: catalase;
GFP: green fluorescent protein;
GORK: Outward-Rectifying K^+^-efflux channel;
HAK5: High-Affinity K^+^ uptake;
HMA: Heavy Metal-Associated;
IRT: Iron-Regulated Transporter and MDA, malondialdehyde;
MIFE: microelectrode ion flux estimation;
NBR1: neighbor of BRCA1;
NRAMP: Natural-Resistance-Associated Macrophage;
POX: peroxidase;
ROS: reactive oxygen species;
RPT5A: 26s proteasome subunit;
SnRK: Snf2-related protein kinase;
TOR: target of rapamycin;
ZIP: Zrt-/Irt-like protein;
YSL3: Yellow Stripe-Like.

---

**Supplemental table S1**. Primers used for quantitative PCR.

**Supplemental figure 1**. **Autophagy flux in response to growing Cd concentrations and time of treatment**. **A**, Fluorescence intensity in leaves disc from Arabidopsis lines expressing the ATG8a-GFP exposed to Cd 50 µM during 1, 3 7 and 10 days. **B** Western blot of α-ATG8 in Arabidopsis wild type, histograms show mean ± SEM of 3 replicates of ATG8 and ATG8-PE (lipidated) (16 kDa and 13 kDa, respectively) from replicates. Values followed by different letters are statistically significant at p<0.05.

**Supplemental Figure 2. Heatmap of ionomic in shoots and roots.** Data were obtained by using Inductively coupled plasma mass spectrometry (ICP-MS) from leaves and roots from Arabidopsis plants treated with Cd 50µM for 3 days. Clustering distance was calculated using correlation, and Ward’s method was applied to generate hierarchical clusters. The color scale represents low values in white and high values in blue colours.

**Supplemental Figure 3. Heatmap of transporters expression**. Data were obtained from qPCR from transporters analysed in this work in shoots and roots from Arabidopsis plants treated with cadmiun 50µM for 3 days. Clustering distance was calculated using correlation, and Ward’s method was applied to generate hierarchical clusters. The color scale represents lower values in white and the highest values in blue color. Letters A (A1 and A2), B (B1 and B2), C and D (D1 and D2) identify each subcluster obtained.

**Supplemental movie S1**. Time laps imaging of autophagosome formation in response to Cd treatment (100 μM; 0-7 d). Seven days-old seedlings of WT (Col-0) and ATG8a-GFP lines were incubated in multiwell plates with MS medium with and without Cd. GFP was imaged at Ex/Em 488/530 (first and third columns) and Chlorophylls at Ex/Em 630/680 (second and fourth columns), used as reference of leaves. Imaging was carried out in a Clariostar

