## Supplemental files for "Deciphering the role of autophagy under Cd toxicity in *Arabidopsis thaliana*"

**Supplemental table S1.** Primers used for quantitative PCR

| Gen | ID | Sequence (5'-3') |
|---|---|---|
| *ATG4a* | AT2G44140 | F: GGCTGCATTGCAACTAGATTT<br>R: GAATCATGCAACCCCAGTTC |
| *ATG4b* | AT3G59950 | F: CTTTCACGTTCCCTCAAAGC<br>R: TTGCAATGGTAAGACGATGTG |
| *ATG5* | AT5G17290 | F: CCTTTGTGCAGAACCCGAAA<br>R: TCCTCTTGATCACTCTGAGACA |
| *ATG7* | AT5G45900 | F: CAATCTCTCAGAAAGATAAGATCAGCCATG<br>R: GAAGCCACCTAGTGAATATAGTCTATGGAC |
| *ATG8a* | AT4G21980 | F: CGATCTTTGGATGACTTTGATG<br>R: TGACGATTAATAAACCCAAAGG |
| *ATG8h* | AT3G06420 | F: CCAAAGCTCTCTTTGTTTTCG<br>R: AAGAACCCGTCTTCTTCCTTG |
| *TOR* | AT1G50030 | F: ATTTCTTCTGCGATCTTCGG<br>R: CAACACTTGGGCAGAGAGAG |
| *HMA2* | AT4G30110 | F: GACAACTACTGCAGCTTCTG<br>R: GCTGTAGCATCTCTTGCAAG |
| *HMA4* | AT2G19110 | F: AGGCCTAGGATCGACATCAAC<br>R: GGTGAATAGGAACACAACTGC |
| *IRT1* | AT4G19690 | F: TTCACTCGGTGGTCATTGGA<br>R: CCGAATGGTGTTGTTACCGC |
| *IRT3* | AT1G60960 | F: GCCCTCACAACCCCGATAG<br>R: GCTCCGACACTGTGAGAATTGA |
| *YSL3* | AT5G53550 | F: ATTGGCCAGGAAACAAGTGTTTGGGT<br>R: GACAAGTCCCGCGACTACACCATTT |
| *Nramp3* | AT2G23150 | F: ATGGTTTTGTGGGTTATGGC<br>R: CTCGAGCTTCCTTATTCCGT |
| *Nramp6* | AT1G15960 | F: GTCAGAGTTCAACCATAA<br>R: GATTATAGCCAAGCTCT |
| *ZIP5* | AT1G05300 | F: CTTCTTGTTGGAGCAGGGGT<br>R: TTGTCAAGAAACGAGAAGGGT |
| *HAK5* | AT4G13420 | F: AAGAGGAACCAAATGCTGA<br>R: GCCCGATGAAGGGACAT |
| *GORK* | AT5G37500 | F: GCTCCATCCGATAG<br>R: CCTCCTTTAATTTAGAAG |
| *TUB4* | AT5G44340 | F: GAGGGAGCCATTGACAACATCTT<br>R: GCAACAGTTCACAGCTATGTTCA |

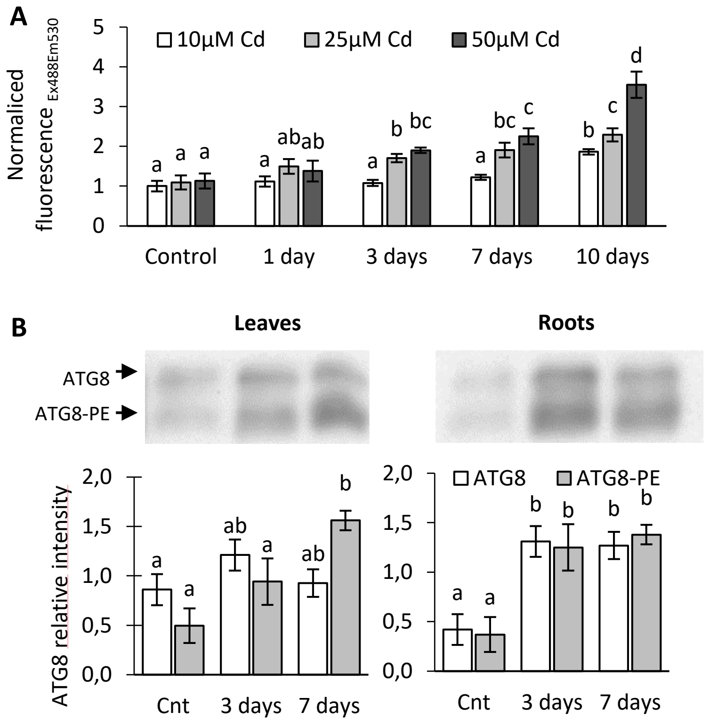


**Supplemental figure 1**. **Autophagy flux in response to growing Cd concentrations and time of treatment**. **A**, Fluorescence intensity in leaves disc from Arabidopsis lines expressing the ATG8a-GFP exposed to Cd 50 µM during 1, 3 7 and 10 days. **B** Western blot of α-ATG8 in Arabidopsis wild type, histograms show mean ± SEM of 3 replicates of ATG8 and ATG8-PE (lipidated) (16 kDa and 13 kDa, respectively) from replicates. Values followed by different letters are statistically significant at $p<0.05$.

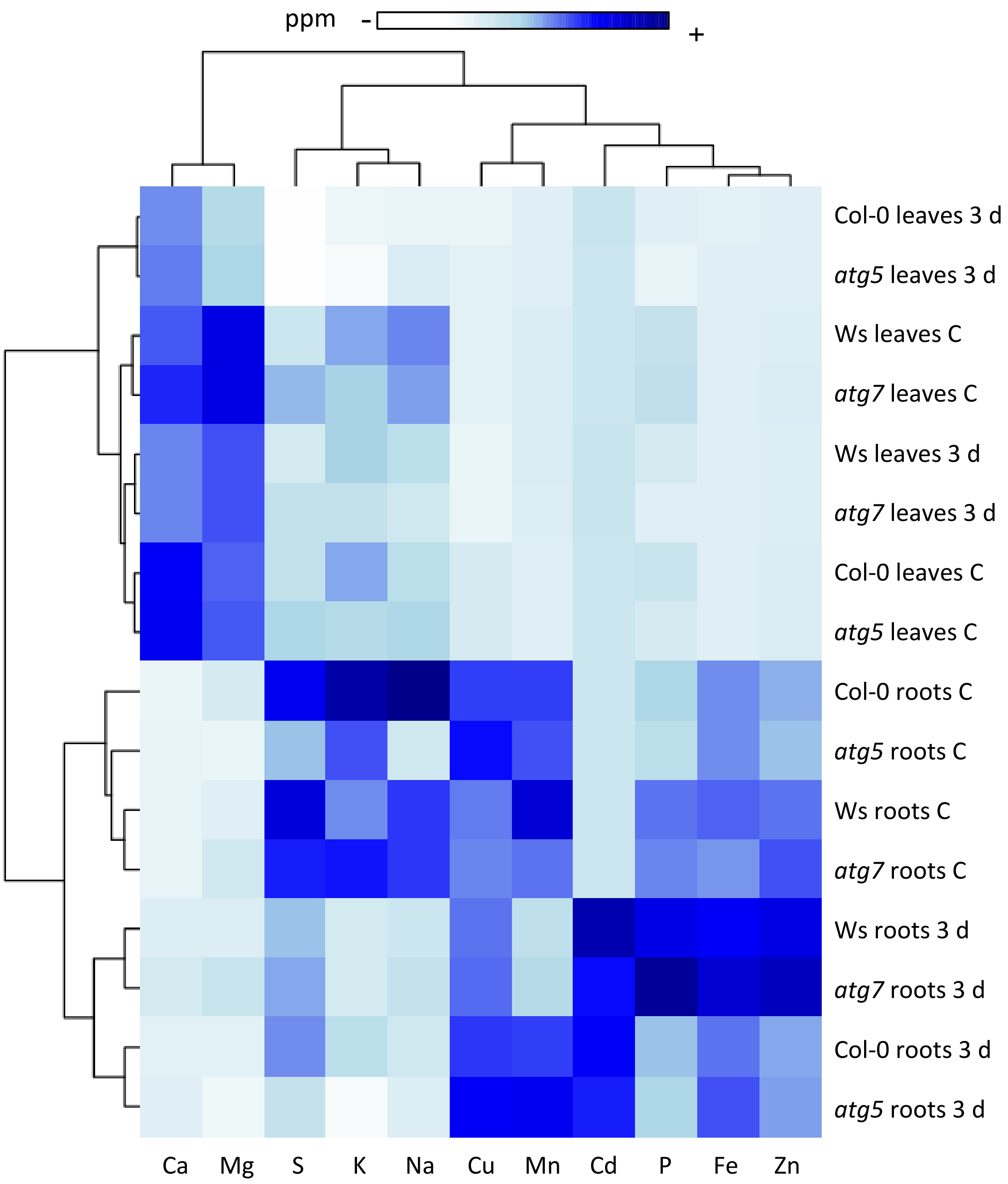


**Supplemental Figure 2. Heatmap of ionomic in shoots and roots.** Data were obtained by using Inductively coupled plasma mass spectrometry (ICP-MS) from leaves and roots from Arabidopsis plants treated with Cd 50µM for 3 days. Clustering distance was calculated using correlation, and Ward's method was applied to generate hierarchical clusters. The color scale represents low values in white and high values in blue colours

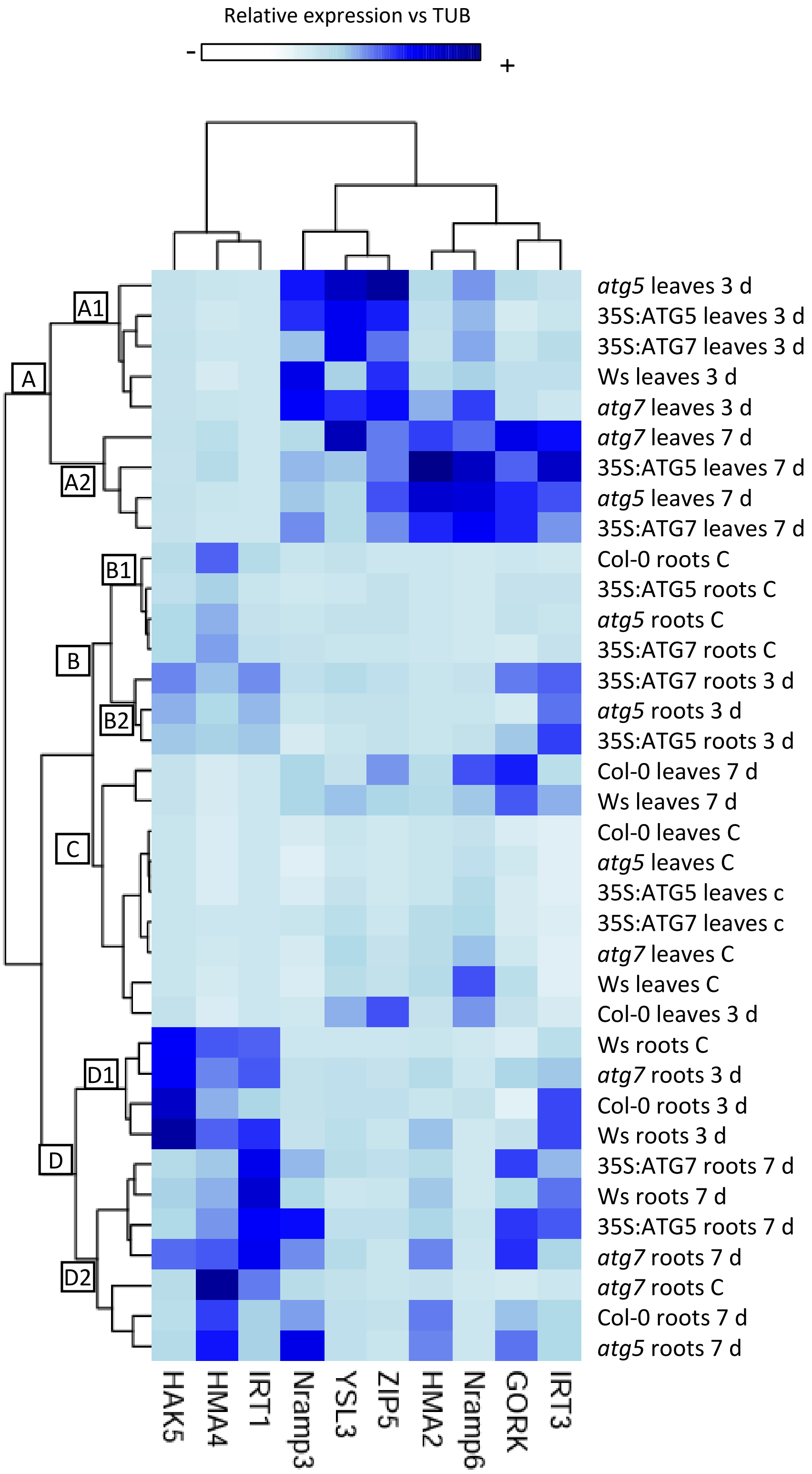


**Supplemental Figure 3**. **Heatmap of transporters expression**. Data were obtained from qPCR from transporters analysed in this work in shoots and roots from Arabidopsis plants treated with cadmiun 50µM for 3 days. Clustering distance was calculated using correlation, and Ward's method was applied to generate hierarchical clusters. The color scale represents lower values in white and the highest values in blue color. Letters A (A1 and A2), B (B1 and B2), C and D (D1 and D2) identify each subcluster obtained.
